# ancIBD - Screening for identity by descent segments in human ancient DNA

**DOI:** 10.1101/2023.03.08.531671

**Authors:** Harald Ringbauer, Yilei Huang, Ali Akbari, Swapan Mallick, Nick Patterson, David Reich

**Author notes:** These authors contributed equally to this work (Harald Ringbauer, Yilei Huang). Corresponding authors –.

## Abstract

Long DNA sequences shared between two individuals, known as Identical by descent (IBD) segments, are a powerful signal for identifying close and distant biological relatives because they only arise when the pair shares a recent common ancestor. Existing methods to call IBD segments between present-day genomes cannot be straightforwardly applied to ancient DNA data (aDNA) due to typically low coverage and high genotyping error rates. We present ancIBD, a method to identify IBD segments for human aDNA data implemented as a Python package. Our approach is based on a Hidden Markov Model, using as input genotype probabilities imputed based on a modern reference panel of genomic variation. Through simulation and downsampling experiments, we demonstrate that ancIBD robustly identifies IBD segments longer than 8 centimorgan for aDNA data with at least either 0.25x average whole-genome sequencing (WGS) coverage depth or at least 1x average depth for in-solution enrichment experiments targeting a widely used aDNA SNP set (‘1240k’). This application range allows us to screen a substantial fraction of the aDNA record for IBD segments and we showcase two downstream applications. First, leveraging the fact that biological relatives up to the sixth degree are expected to share multiple long IBD segments, we identify relatives between 10,156 ancient Eurasian individuals and document evidence of long-distance migration, for example by identifying a pair of two approximately fifth-degree relatives who were buried 1410km apart in Central Asia 5000 years ago. Second, by applying ancIBD, we reveal new details regarding the spread of ancestry related to Steppe pastoralists into Europe starting 5000 years ago. We find that the first individuals in Central and Northern Europe carrying high amounts of Steppe-ancestry, associated with the Corded Ware culture, share high rates of long IBD (12-25 cM) with Yamnaya herders of the Pontic-Caspian steppe, signaling a strong bottleneck and a recent biological connection on the order of only few hundred years, providing evidence that the Yamnaya themselves are a main source of Steppe ancestry in Corded Ware people. We also detect elevated sharing of long IBD segments between Corded Ware individuals and people associated with the Globular Amphora culture (GAC) from Poland and Ukraine, who were Copper Age farmers not yet carrying Steppe-like ancestry. These IBD links appear for all Corded Ware groups in our analysis, indicating that individuals related to GAC contexts must have had a major demographic impact early on in the genetic admixtures giving rise to various Corded Ware groups across Europe. These results show that detecting IBD segments in aDNA can generate new insights both on a small scale, relevant to understanding the life stories of people, and on the macroscale, relevant to large-scale cultural-historical events.

## Introduction

Some pairs of individuals share long, nearly identical genomic segments, so-called Identity-By-Descent (IBD) segments, that must be co-inherited from a recent common ancestor because they are rapidly broken apart by recombination events. Recombination in ancestors marks the end of IBD segments, thus the probability that an IBD segment of a given map length *l* persists for *t* generations decays exponentially as exp(–2*lt*). Consequently, long IBD segments provide an ideal signal to probe recent genealogical connections, and have been used as a distinctive signal for a range of downstream applications such as identifying biological relatives or inferring recent demography [e.g. Palamara and Pe’er, 2013, Ralph and Coop, 2013, Ringbauer et al., 2017].

Several existing methods identify IBD segments for single nucleotide polymorphism (SNP) array or whole genome sequence data [e.g. Gusev et al., 2009, Browning and Browning, 2011, Zhou et al., 2020], but they require high-quality genotype data. For ancient DNA, however, in most cases, it is not possible to confidently call diploid genotypes or confident genotype likelihoods. Typically, aDNA data is of far too low genomic coverage (<5× average coverage per site) and moreover, error rates are comparably high because of degraded and short DNA molecules. The promise of haplotype sharing signals in aDNA research has been acknowledged [Racimo et al., 2020], but so far only a few exceptional applications to comparably high-quality aDNA data have been published [see e.g. Sikora et al., 2017, Ferrando-Bernal et al., 2020]. First efforts to identify IBD based on imputed data have been fruitful [Kivisild et al., 2021, Allentoft et al., 2022, Ariano et al., 2022, Severson et al., 2022a], but those earlier works require higher coverage not routinely available for aDNA. Importantly, they do not include a systematic evaluation of the IBD calling pipelines but performance when screening for IBD is expected to decay for short segments and low-coverage data. Practical downstream applications, such as demographic modeling, require information about power, length biases, and false positive rates - either to account directly for these error processes or to identify thresholds of data quality at which these error processes can be safely ignored.

Here we present and systematically evaluate ancIBD, a method to detect IBD segments in human aDNA data. In brief, ancIBD starts from phased genotype likelihoods imputed by GLIMPSE [Rubinacci et al., 2021], which are then screened using a Hidden Markov Model (HMM) to infer IBD blocks (Fig. 1). We identified default parameters that optimize performance when using a set of ca. 1.1 million autosomal SNPs that have been targeted for in-solution enrichment experiments that have produced more than 70% of genome-wide human ancient DNA data sets to date [Fu et al., 2014, 2015, Rohland et al., 2022]. Our tests show that ancIBD robustly identifies IBD longer than 8 centimorgan in aDNA data for SNP capture aDNA with at least 1x average coverage depth (on target), and for whole-genome sequencing (WGS) even as low as 0.25x average genomic coverage.

**Figure 1:**
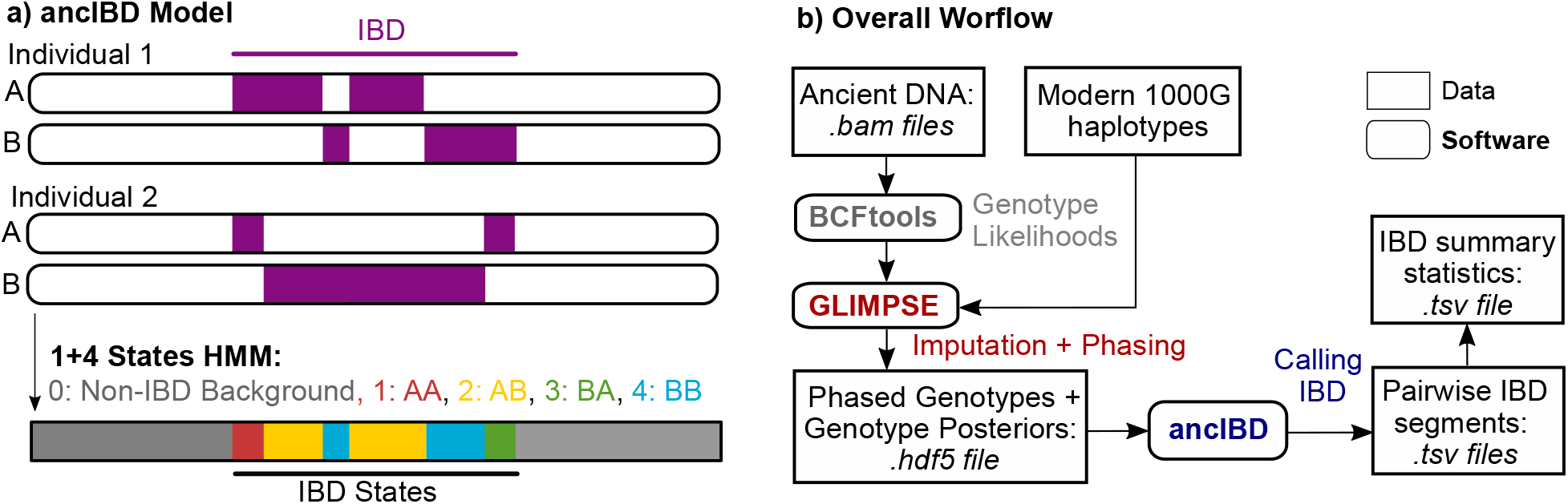
Overview of the ancIBD algorithm. **a)** Sketch of the ancIBD Hidden Markov Model. The ancIBD HMM has five states: one background state of no allele sharing, and four states modeling the four possible IBD sharing states between two phased diploid genomes. We model phase switch errors within a true IBD segment as a transition between the four IBD states. **b)** Visualization of the full pipeline to call IBD. First, ancient DNA data is imputed and phased using published software (GLIMPSE) and a panel of modern reference haplotypes, and then our HMM (ancIBD) is applied to screen for IBD. The final output consists of two .tsv tables, one listing all inferred IBD segments and one listing IBD summary statistics for each pair of individuals.

## Methods

Our method consists of two computational steps (Fig. 1b): First, the aDNA data is computationally imputed and phased. Second, we apply a custom HMM to output IBD.

For the first step, we use existing software that has been shown to work well for low coverage data, GLIMPSE [Rubinacci et al., 2021], which we apply to aligned sequence data (in .bam format) to impute genotype likelihoods at the 1240k sites, using haplotypes in the 1000 Genome project as reference panel (see Supp. Note S3). Previous evaluation on aDNA data showed that imputed genotypes for common variants, that are highly informative about IBD sharing, are of relatively good quality down to mean coverage depth as low as 0.5-1.0x[Hui et al., 2020, da Mota et al., 2022].

### The Hidden Markov Model

The new HMM makes use of the imputed genotype probabilities and phase information output by GLIMPSE and, for each pair of samples, runs a forward-backward algorithm [Durbin et al., 1998] to calculate the posterior probabilities of being in an IBD state at each marker (Fig. 1). These probabilities are then post-processed to call IBD segments. In the following sections, we describe this HMM (Fig. 1a) in detail, in particular its states, the model for emission and transition probabilities, the calling of IBD segments and postprocessing, and its implementation.

Throughout, we assume biallelic variants and denote the two individuals we screen for IBD as 1 and 2, and their phased haplotypes as (1*A*, 1*B*) and (2*A*, 2*B*). The HMM screens each of the 22 autosomal chromosomes from beginning to end independently, thus it suffices to describe the HMM applied to one chromosome.

#### Hidden States

Our Hidden Markov model has five hidden states *s* = 0,1…4. The first state *s* = 0 encodes a non-IBD state, while the four states *s* = 1,2,3,4 encode the four possibilities (1A/2A, 1A/2B, 1B/2A, 1B/2B) of sharing an IBD allele between the haplotypes of two diploid genomes (1A,1B) and (2A,2B) (Fig. 1a). We note that we do not model IBD sharing beyond a single pair of haplotypes (where both pairs of or more than three haplotypes share a recent common ancestor). These cases occur only rarely in practice [Chiang et al., 2016] and our goal here is to identify long tracts of IBD.

#### Transition Probabilities

To calculate the 5 × 5 transition probabilities *T* to change states from one to the following loci, denoted by *l* and *l* + 1, we make use of the genetic map distances obtained from a linkage map, i.e. a map of the position using Morgans as the unit of length (one Morgan is the genomic map span over which the average number of recombinations in a single generation is 1).

As in [Ringbauer et al., 2021], we specify the transition probabilities via a 5 × 5 infinitesimal transition rate matrix *Q*, from which each transition probability matrix *A*_*l*→*l*+1_ is obtained via matrix exponentiation using the genetic distance *r_l_* between loci *l* and *l* +1

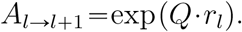

Here, *Q* is defined by the following three rate parameters: The rate to jump from the non-IBD state into any of the four IBD states (IBDin), the rate to jump from any of the IBD states to the non-IBD states (IBDout), and the rate to jump from any of the IBD states to another one (IBDswitch):

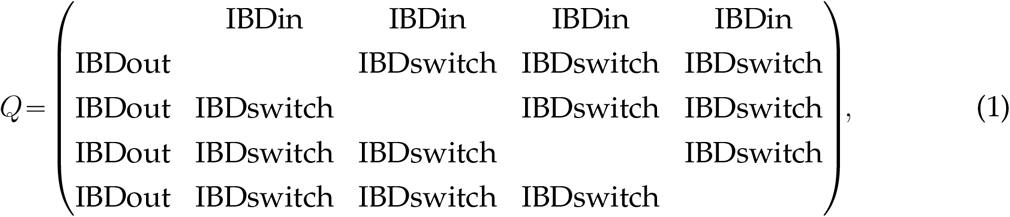

where the diagonal elements are defined as *Q_ii_* =∑_*j*≠*i*_ *Q_ij_* such that the rows of *Q* sum to zero as required for a transition rate matrix. The rate IBDswitch models phasing errors, as a transition from one IBD state to another means that a different haplotype pair is shared. We note that the probability of the IBD state jumping from 1A/2A to 1B/2B would require phase switch errors to occur in both individuals at the same genomic location, which is highly unlikely; however, we set the transition matrix between all four IBD states symmetric as this allowed us to implement a substantial computational speed up.

### Emission Probabilities

#### Single Locus Emission Probabilities

To define the emission model of the HMM, we need to specify *P*(*D*|*s*), the likelihood of the genetic data for the five HMM states *s* =0,1,⋯,4 at one locus. Throughout, we denote reference and alternative alleles as 0 and 1, respectively, and the corresponding genotype as *g*∈{0,1}. The observed data *D* of our emission model will be the haploid dosage, which is the probability of a phased haplotype carrying an alternative allele, here denoted for each haplotype *h* as

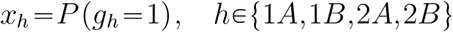

First, we explain how we approximate the two haploid dosages for a single imputed diploid individual 1. We have to use an approximation as GLIMPSE only outputs the most likely phased diploid genotype *GT*∈{0|0,011,110,111} as well as three posterior genotype probabilities *GP* for each of the unphased diploid genotypes, denoted by the number of alternative alleles as 0,1,2. We first approximate the posterior probabilities for the four phased states, here denoted as *P*_00_, *P*_01_, *P*_10_, *P*_11_. The two homozygote probabilities *P*_00_ and *P*_11_ are obtained trivially from the corresponding unphased genotype probabilities *GP*, as no phase information is required for homozygotes. To obtain probabilities of the two phased heterozygotes states, *P*_01_ and *P*_10_, we use a simple approximation. Let *p*_0_, *p*_1_, *p*_2_ denote the posterior probability for each of the three possible diploid genotypes. If the maximum-likelihood unphased genotype is heterozygote, i.e. *max*(*p*_0_, *p*_1_, *p*_2_) = *p*_1_, we set *P*_01_ = *p*_1_, *P*_10_ = 0 if *GT*=0|1 and *P*_01_ = 0, *P*_10_ = *p*_1_ if *GT* = 1|0. If the maximum-likelihood unphased genotype is a homozygote, i. e. *max*(*p*_0_, *p*_1_, *p*_2_) = *p*_0_ or *p*_2_, and thus there is no phase information for the heterozygote genotype available, we set *P*_01_ = *P*_10_ = *p*_1_{2. Having obtained the four probabilities for the possible phased genotypes, we can calculate the two haploid dosages as:

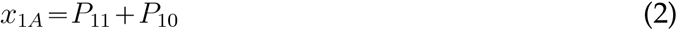

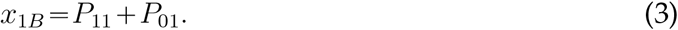

When calling IBD segments between two individuals 1 and 2, we use this approach to obtain all four haploid dosages and denote them for haplotypes 1*A*, 1*B*, 2*A*, 2*B* as (*x*_1*A*_, *x*_1*B*_, *x*_2*A*_, *x*_2*B*_).

Setting those four haploid dosages as the observed data *D* = (*x*_1*A*_, *x*_1*B*_, *x*_2*A*_, *x*_2*B*_) at one locus, we can now calculate the likelihood *P*(*D*|*s*) for each of the five HMM states *s* = 0,1,⋯,4. We start by summing over all possible unobserved latent phased genotypes **g**= (*g*_1*A*_, *g*_1*B*_, *g*_2*A*_, *g*_2*B*_), yielding in total 16 possible combinations of reference and alternative alleles, denoted together as 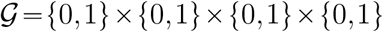:

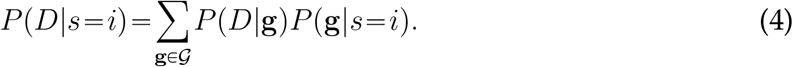

For the term *P*(*D* |**g**), applying Bayes rule yields:

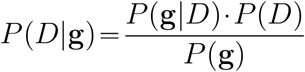

*P*(*D*) remains a constant factor across all states, which can be ignored because posterior probabilities of an HMM remain invariant to constant factors in the likelihood. We arrive at:

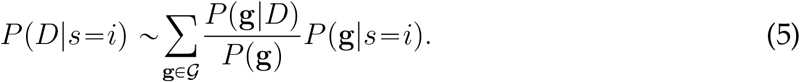

We now approximate the three quantities on the right-hand side of eq. 5 for a given set of genotypes **g**.

First, assuming Hardy-Weinberg equilibrium, *P*(**g**) is calculated as the product of the four corresponding allele frequencies of (either *p* or 1 – *p* depending on the respective allele in **g** being 0 or 1). In practice, we obtain *p* from the allele frequencies in the reference panel.

Second, we approximate *P*(**g**|*D*) as the product of the four probabilities of each of the haplotypes (1A, 1B) and (2A, 2B) being reference or alternative. We assume that diploid genotype probabilities can be approximated as products of the respective haploid dosages, which we empirically verified on GLIMPSE imputed data (Fig. S14). Using the haploid dosages (*x*_1*A*_, *x*_1*B*_, *x*_2*A*_, *x*_2*B*_) as calculated above yields:

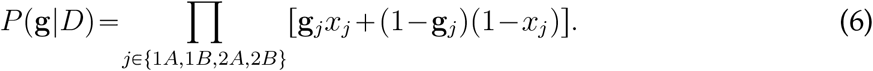

Third, to approximate *P*(**g** | *s* = *i*) we again assume Hardy-Weinberg probabilities which yield a product of factors *p* or 1 – *p* (listed in Tab. S1). For the four IBD states, the two shared alleles constitute one shared draw. Consequently, there are only three instead of four independent factors, and genotype combinations **g** where the shared genotype would be different have 0 probability.

Plugging these three approximations into eq. 5 now give *P*(*D*|*s*) for each state *s* = 0,1… 4.

For the background state (*s*=0) we have *P*(*g*) = *P*(*g* | *s* = 0) and thus these factors cancel out in eq. 5. Using that ∑_g_*P*(**g**|*D*) = 1, we arrive at:

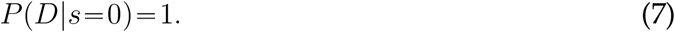

The four IBD states (*s* = 1,2,3,4) are calculated analogously with a simple rearrangement of the haplotype order. Thus, it suffices to describe *s* = 1, the state where the two first phased genotypes, 1A and 2A, are identical. For the two non-shared alleles the Hardy-Weinberg factors cancel out as in *s*=0. After some rearranging (see Supp. Note S1), we obtain:

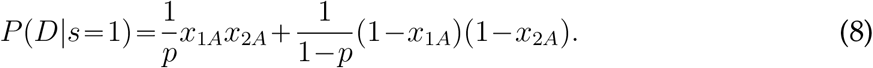

### Post-processing: Calling IBD segments

To call IBD segments, we use the posterior probability of being in the IBD states obtained via the standard HMM forward-backward algorithm [Bishop, 2006], which takes as input the transition rates (eq. 1) and emission probabilities (eq. 7 and eq. 8). Our method then screens for consecutive markers where the posterior probability of being in the non-IBD state *h*=0 remains below a pre-specified threshold. We determine the start of an inferred IBD segment by locating the first SNP whose posterior decreases below the threshold and the end by the first SNP whose posterior rises above the threshold. For each such genomic region longer than a pre-specified minimum length cutoff, one IBD segment is recorded.

#### Merging Spurious gaps between IBD

A post-processing step commonly applied when detecting IBD is to merge two closely neighboring IBD segments [Browning and Browning, 2011, Ralph and Coop, 2013]. This step aims to remove spurious gaps within one true IBD segment, which can appear due to low density of SNPs or due to sporadic genotyping errors. The rationale is that under most demographic scenarios sharing of long IBD is very rare and thus two IBD segments are unlikely to occur next to each other by chance [Chiang et al., 2016]). Removing artificial gaps is important for determining the length of an IBD segment, and therefore in particular for downstream methods that use the lengths of IBD segments as a recombination clock. In our implementation, we merge all gaps where both IBD are longer than a threshold length and separated by a gap of a maximum length.

#### Applying genomic masks

To exclude regions prone to false positive IBD, we designed a series of genomic masks by examining rates of IBD segments across the genome when inferring IBD in a large set of empirical aDNA data. As described in Supp. Note S4, Fig.S4), about 8% (a total of 13 regions) of the genome is masked, with most masked regions involving centromeres and telomeres.

### Setting default parameters of ancIBD

To avoid batch effects between WGS and 1240k data, ancIBD generally takes as input imputed 1240k sites. In the following, we describe how we choose the default parameters of ancIBD to work well for imputed genotype likelihoods at this 1240k SNP set.

First, we simulated a dataset including ground-truth IBD sharing by using haplotypes in the 1000 Genome project panel. We simulated chromosome 3 by stitching together short haplotypes 0.25 cM long copied from individuals labeled as TSI (Tuscany, Italy) in the 1000 Genomes Project data [Consortium et al., 2015] and then copied IBD segments of various lengths (4,8,12,16,20 cM) into 100 pairs of mosaic genomes (described in detail in Fig. S1, Supp. Note S2). This approach, following Ralph and Coop [2013], yields a set of diploid genotype data with exactly known IBD. Such a haplotype mosaic removes long IBD segments in the 1000 Genome data while also maintaining most of the local haplotype structure. To obtain data typical for aDNA sequencing, we matched genotyping errors and probabilities observed within downsampled high-coverage empirical aDNA data and added phase switch errors (as described in Supp. Note S2).

We then applied ancIBD for a range of parameter combinations and recorded performance statistics (Supp. Tab. 1D and 1E). The final parameters that we set as default values (listed in Tab. 1) are chosen to work well for a broad range of coverages and IBD lengths. Throughout this work, we use these settings, but in our implementation, each parameter can be changed to a non-default value by the user.

**Table 1:** Parameters of ancIBD HMM and default values. All parameters that can be set in our implementation. The default values are chosen to work well (low FP, high power, little length bias and variation) for a broad range of WGS and 1240k aDNA data (see Supp. Tab. 1D and 1E).

| Parameter | Value | Description [Unit] |
| --- | --- | --- |
| **HMM** |  |  |
| ibd_in | 1 | Transition rate out of non-IBD state [per Morgan] |
| ibd_out | 10 | Transition rate out of IBD states [per Morgan] |
| ibd_jump | 400 | Transition rate between IBD states [per Morgan] |
| min_error | 0.001 | Cap Min. Probability of diploid genotype Error [rate] |
| p | Sample/1000G | Allele Frequencies (AF) |
| **Post-Processing** |  |  |
| cutoff_post | 0.99 | Minimum Posterior on IBD state for IBD segment call |
| min_cm | 2 | Minimum Length IBD segment to call [centimorgan] |
| snp_cm | 220 | Minimum SNP density in IBD [per centimorgan] |
| max_gap | 0.0075 | Maximum Gap to merge [Morgan] |

### Implementation and Runtime

We implemented a number of computational speed-ups to improve the run-time of our algorithm. First, the forward-backward algorithm is coded in the *Cython* module to make use of the increased speed of a pre-compiled C function within our overall Python implementation. Second, our algorithm utilizes a re-scaled version of the forward-backward algorithm [Bishop, 2006] which avoids computing logarithms of sums that would be computationally substantially more expensive than products and additions. Finally, we make use of the symmetry of the four IBD states. As the transition probabilities between those are fully symmetric, we can reduce the transition matrix from a 5×5 to a 3×3 matrix by collapsing the three other IBD states into a single “other IBD” state. After the exponentiation of the 3×3 matrix, the original 5×5 transition matrix is reconstructed by dividing up the jump rates using the original symmetry.

We transform the VCF output of GLIMPSE to an HDF5 file, a data format that allows efficient partial access to data [The HDF Group, 1997-2023], e.g. we can effectively load data for any subset of individuals.

The average runtime of ancIBD for a pair of imputed individuals on all 22 autosomes is about 25 seconds when using a single Intel Xeon E5-2697 v3 CPU with 2.60GHz (Fig. E1). As the number of pairs in a sample of *n* individuals grows as *n*(*n*–1)/2, the runtime scales quadratically when screening all pairs of samples for IBD (Fig. E1). However, we note that due to the speed of a HMM forward-backward algorithm with 5 states requiring only a few multiplications and additions per locus, a large fraction of runtime per pair is due to loading the data (Fig. E1). Thus, an efficient strategy is to load a set of individuals into memory jointly, as then the loading time scales only linearly with the number of samples. This strategy, implemented in ancIBD, leads to hugely improved runtime per pair of samples in cases where many samples are loaded into memory and screened for pairwise IBD (Fig. E1). We observed that for batches of size 50 samples and when screening all 50 · 49/2 = 1225 pairs for IBD, the average runtime of ancIBD per imputed pair for all 22 chromosomes reduces to ca. 0.75 seconds. The asymptotic limit per sample pair, which is the runtime of the HMM and post-processing, is about 0.35 seconds on our architecture.

## Results

We performed two sets of experiments to evaluate the quality of IBD calls of ancIBD at various coverages. First, we copied IBD segments of known length into pairs of genomes (as described in the section “Setting Default Parameters of ancIBD). Second, we downsampled high-coverage empirical aDNA data. We describe both methods and our results in the following section.

### Performance on copied-in IBD segments

When applying ancIBD to the simulated data with copied-in IBD, we observed that the inferred IBD segments remain accurate and that their length distribution peaks around the true value for WGS data down to about 0.25x coverage, and for 1240k capture data down to 1x coverage at 1240k sites (Fig. 2). We found that ancIBD on average overestimates the length of IBD segments, but within the recommended coverage cutoff the length errors remain within ~1 cM (see Tab. E2, Tab. E3).

**Figure 2:**
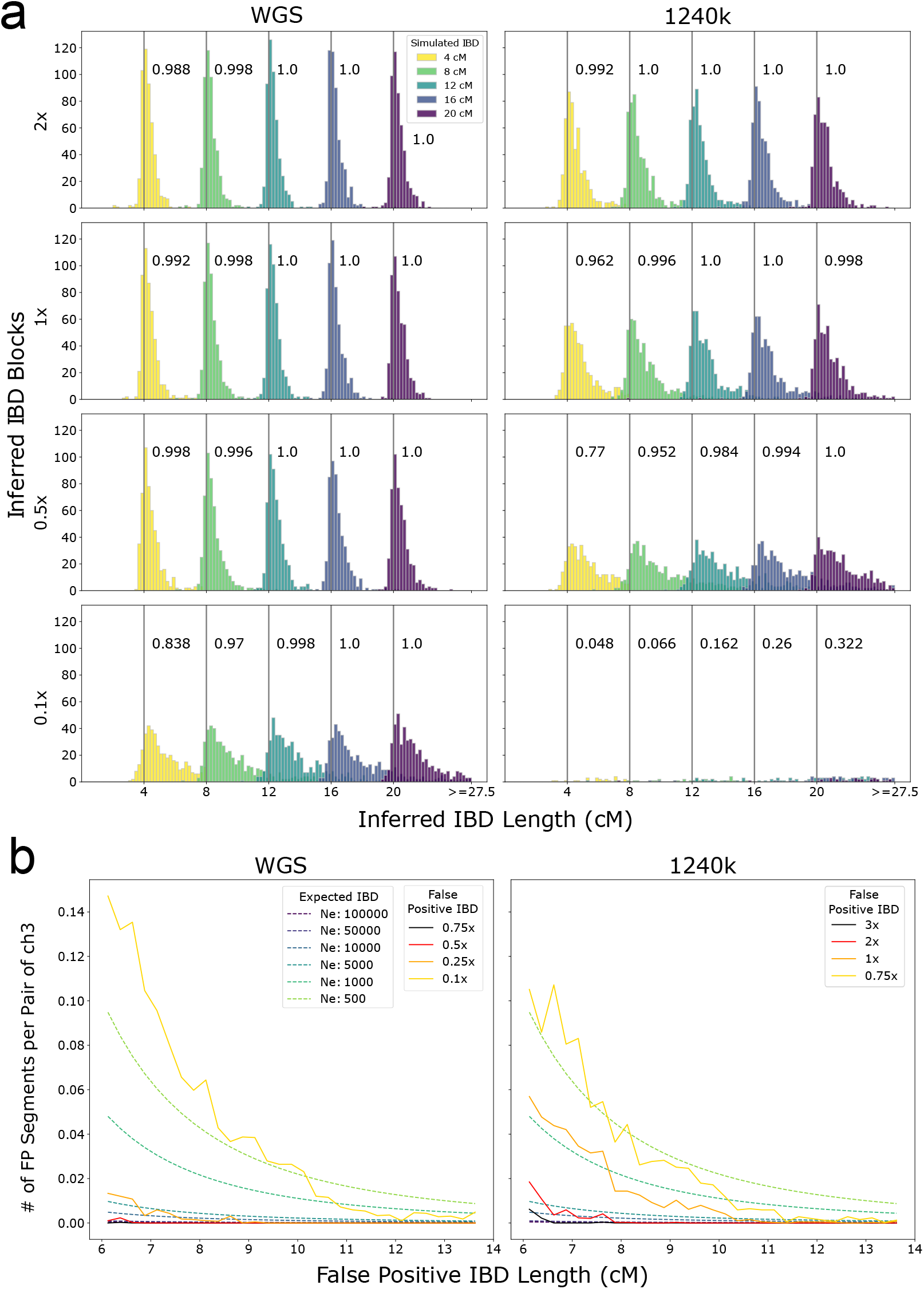
Performance of ancIBD on simulated IBD segments. a: Power and segment length errors. We copied in IBD segments of lengths 4,8,12,16,20 cM into synthetic diploid samples. We simulated shotgun-like and 1240k-like data (as described in Supp. Note S2) and visualize false positive, power, and length bias for 1x, 0.5x, and 0.25x coverage (rows). For each parameter set and IBD length, we simulated 500 replicates of pairs of chromosome 3, each pair with one copied-in IBD segment. The power (or recall) of detecting IBD segments of each simulated length is indicated in the text next to the corresponding gray vertical bar. **b: False positive rate**. We downsampled high-quality empirical aDNA data without IBD segments (Supp. Tab. 1F) to establish false positive rates of IBD segments for various coverage and IBD lengths (described in Supp. Note S5). We visualized expected IBD sharing assuming various constant population sizes to contextualize the false positive rates.

### Performance on downsampled ancient DNA data

To assess performance on downsampled empirical aDNA data, we used four high-coverage genomes of ancient individuals, all ca. 5000 years old and associated with the Afanasievo culture (Supp. Note S4). When comparing the IBD calls in the downsampled data to the IBD calls of the original high-coverage data, we found that WGS substantially outperforms 1240k data of the same coverage. For long IBD segments (>10cM) that are particularly informative when detecting relatives, ancIBD achieves high precision and recall (>90%) for all coverages tested here (WGS data: 0.1x-5x, 1240k data: 0.5x-2x). For intermediate range segments (8-10cM), ancIBD maintains reasonable recall (~80%) at all coverages while having less than 80% precision at 0.5x for 1240k data. Overall, ancIBD yields accurate IBD calling (~90% or higher precision) at >0.25x wgs and >1x 1240k data (Fig. S).

### Comparing to other methods

Several recent publications have applied methods designed to detect IBD in high-quality present-day data on imputed aDNA data (e.g. using GLIMPSE) [Kivisild et al., 2021, Allentoft et al., 2022]. To compare the performance of ancIBD to such methods, we used the downsampled empirical ancient aDNA data described above.

Software to call IBD can be classified into two categories, ones that require prior phasing and ones that use unphased data as input. The former search for long, identical haplotypes, while the latter mostly utilize, directly or implicitly, the signal of “opposing homozygotes” (two samples being homozygous for different alleles), which are lacking in IBD segments.

In preliminary tests, we found that methods that require phasing information have very low power to detect IBD in imputed aDNA data, potentially because of high switch error rates in imputed ancient genomes [da Mota et al., 2022], which is an order of magnitude higher than what is attainable for phasing biobank scale modern data [Delaneau et al., 2019].

Therefore, we focus our detailed comparison on two methods that do not require phasing information, IBIS [Seidman et al., 2020] and IBDseq [Browning and Browning, 2013a]. IBIS detects IBD segments by screening for genomic regions with few opposing homozygotes. Our results on downsampled aDNA data show that this method tends to maintain higher precision at the expense of a lower recall, particularly at lower coverages. Despite keeping precision at >90%, for segments >8cM, I BIS’s recall drops to ~ 50% for ~1x 1240k data (Fig. S3).

IBDseq is originally designed for whole genome sequencing data. It works by computing likelihood ratios of IBD and non-IBD states for each marker and then identifies IBD segments by searching for regions with high cumulative scores. Our results on downsampled empirical ancient aDNA data indicate that IBDseq’s precision and recall drop substantially at lower coverages. With less than 50% precision for ~1x 1240k data, IBDseq would perform poorly for most aDNA samples (Fig. S10, Fig. S11).

### Detecting close and distant relatives with ancIBD

To showcase the utility of IBD segments to detect biological relatives, we applied ancIBD to a large set of aDNA data of ancient Eurasians (a superset of the AADR dataset containing published and unpublished aDNA data, see Data Availability). After filtering to all individuals with geographic coordinates in Eurasia with dates within the last 45,000 years, and sufficient genomic coverage for robust IBD calling we are left with 10,156 unique ancient individuals (4,274 of which are published, Supp. Tab. 1A). As coverage cutoff, we used at least 600,000 1240k SNPs covered for 1240k data, and at least 200,000 1240k SNPs covered for whole-genome-sequencing aDNA data. This metric was chosen as the number of 1240k SNPs covered is readily available in the AADR database for all samples. This cutoff corresponds to our recommended minimum coverage cutoff for ancIBD described above, as the relationship between average coverage and the number of SNPs covered is broadly monotonic (see Fig. S13). We then screened each of the 51,567,090 pairs of ancient genomes with ancIBD. To optimize runtime we grouped the 10,156 samples into batches of 400 individuals - and then ran all sample pairs between pairs of batches (this approach is implemented in the in ancIBD software package). For each pair with detected IBD, we collected IBD statistics into a summary table (see Supp. Tab. 1B for pairs of published individuals).

We find that the pattern of IBD sharing closely mirrors simulated IBD sharing between various degrees of relatives (using the software PEDSIM [Caballero et al., 2019]) when plotting the total sum and the total count of IBD segments longer than 12 cM (Fig. 3), but we note that in the empirical dataset slightly less IBD segments than simulated one are identified, likely due to the filtering of IBD segments with too low SNP density. A first-degree relative cluster becomes apparent, with a parent-offspring cluster (where the whole genome is in IBD), and a full-sibling cluster. The parent-offspring cluster in the simulated IBD dataset consists of one point, as expected because parent and offspring share each of the 22 chromosomes fully IBD. In the inferred IBD dataset, the apparent parent-offspring cluster is spread out more widely, including also individuals with a higher number of IBD segments - the reason for this is that sporadically very long IBD are broken up by artificial gaps, and if they are too big they are not merged by the default gap merging of ancIBD. Overall this effect remains modest, and in the parent-offspring cluster the total number of inferred IBD segments is in most cases only slightly elevated beyond the expected 22.

**Figure 3:**
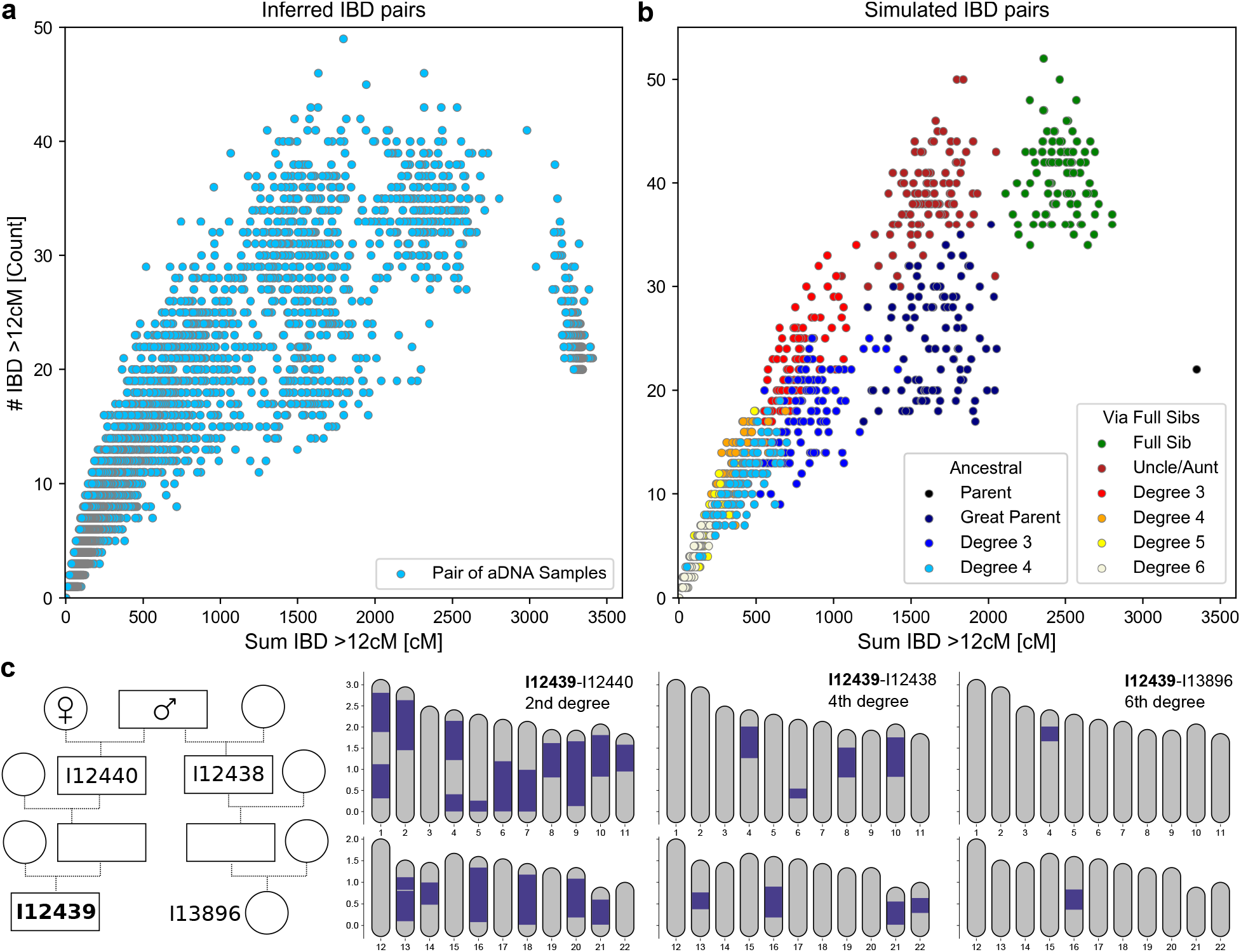
Inferring biological relatives in the ancient DNA record using long IBD inferred with ancIBD. **a**: Inferred IBD among pairs of 10,156 ancient Eurasian individuals. The plot visualizes both the count (y-axis) as well as the summed length (x-axis) of all IBD at least 12 cM long. **b**: Simulated IBD among pairs of relatives. For each relative class, we simulated 100 replicates using the software PEDSIM [Caballero et al., 2019], as described in Supp. Note S6. As in **a** we depict the summed length and the count of all IBD at least 12 cM long. **c**: Inferred IBD among four ancient English Neolithic individuals, who lived about 5,700 years ago and were entombed at Hazleton North long cairn. A full pedigree was previously reconstructed using first and second-degree relatives inferred using pairwise SNP matching rates [Fowler et al., 2022]. We depict all IBD at least 12 cM long. The four individuals were genotyped using 1240k aDNA capture (I12438: 3.7x average coverage on target, I12440: 2.1x, I13896:1.1x, I12439: 6.7x)

Further, we observe two clear second-degree relative clusters that correspond to biological great-parent grandchildren and aunt/uncle-niece/nephew relations ships. Half-siblings are expected to form a gradient between these two clusters, with their average position depending on whether the shared parent is maternal (on average more but shorter shared segments) or paternal (fewer but longer shared segments) [Caballero et al., 2019].

In the simulated data, IBD clusters for 3rd-degree and more distant relatives increasingly overlap, and the empirical IBD distribution follows this gradient. Due to this biological variation in genetic relatedness, it is not possible to uniquely assign individuals to specific relative clusters beyond third-degree relatives even if the exact IBD is known. However, these pairs with multiple long shared segments still unambiguously indicate very recent biological relatedness. The majority of biological relatives up to the sixth degree will share two or more long IBD segments [e.g. Caballero et al., 2019]. In most human populations pairs of biologically unrelated (i.e. related at most by tenth degree) individuals share only very sporadically single IBD segments [Palamara et al., 2012, Carmi et al., 2013, Ringbauer et al., 2021]. Thus, sharing of multiple long IBD segments provides a distinct signal for identifying close genealogical relationships that we can detect with ancIBD. For instance, we identified two long IBD in a sixth-degree relative from Neolithic Britain, a relationship that was previously reconstructed from a pedigree of first-degree and second-degree relatives identified using average pairwise genotype mismatch rates [Fowler et al., 2022].

### Applying ancIBD to Copper and Bronze Age Western Eurasians reveals recent links

Because recombination acts as a rapid clock (the probability of an IBD segment of length l centimorgan persisting for *t* generations declines quickly as exp(–*t*\**l*/50)), the rate of sporadic sharing of IBD segments can probe biological connections between groups of individuals only a few hundred years deep [e.g. for modern Europeans Ralph and Coop, 2013]. To showcase how detecting IBD segments with ancIBD can reveal such connections between ancient individuals, we applied our method to a set of previously published ancient West Eurasian aDNA data dating to the Late Eneolithic and Early Bronze Age (Supp. Tab. 1C). This period, from 3000 to 2000 BCE, was characterized by major gene flow events, where ‘Steppe-related’ ancestry had a substantial genetic impact throughout Europe [e.g. Haak et al., 2015, Allentoft et al., 2015], leading to widespread genetic admixtures as far west as Britain [Olalde et al., 2018] and Iberia [Olalde et al., 2019]. Applying ancIBD to the relevant published aDNA record of 304 ancient Western Eurasians organized into 24 archaeological groups (Supp. Tab. 1C), we find several intriguing links. Many of those connections were previously hypothesized and suggested by admixture tests; however, sharing of long IBD segments now provides definitive evidence for recent co-ancestry and biological interactions, tethering groups together closely in time.

We found that several nomadic Steppe groups associated with the Yamnaya culture that dates to around 3000 BCE share relatively large amounts of IBD with each other (Fig. 5). Notably, this IBD cluster includes also individuals associated with the Afanasievo culture near the Central Asian Altai mountains several thousands of kilometers east. This signal of IBD sharing confirms the previous archaeological hypothesis that Afanasievo and Yamnaya are closely linked despite the vast geographic distance from Eastern Europe to Central Asia [e.g. Anthony, 2010]. A genetic link has already been evident from genomic similarity and Y haplogroups [Allentoft et al., 2015, Narasimhan et al., 2019]; however, the time depth of this connection remained unclear. We find IBD signals across all length scales, importantly including several shared IBD segments even longer than 20 cM (Fig. E2). Such long IBD links must be recent as recombination ends an IBD segment ca. 20 cM long on average every five meiosis. This IBD sharing signal therefore clearly indicates that ancient individuals from Afanasievo contexts descend from people who migrated at most a few hundred years earlier across vast distances of the Eurasian Steppe.

Increased individual mobility in Eneolithic and Early Bronze Age Eurasian Steppe groups is also reflected in a pair of individuals associated with the Afanasievo culture that were buried 1410 km apart, one in present-day Central Mongolia and one in Southern Russia, who share several long IBD segments (Fig. 4). We identified four IBD segments 20-40 cM long, a distinctive signal of close biological relatedness typical of ca. 5th-degree relatives (Fig. 4). Previous work showed that both individuals have a genetic profile typical for Afanasievo individuals, and here this close biological link demonstrates that at least one individual in the chain of relatives between them must have traveled several hundreds of kilometers in their lifetime.

**Figure 4:**
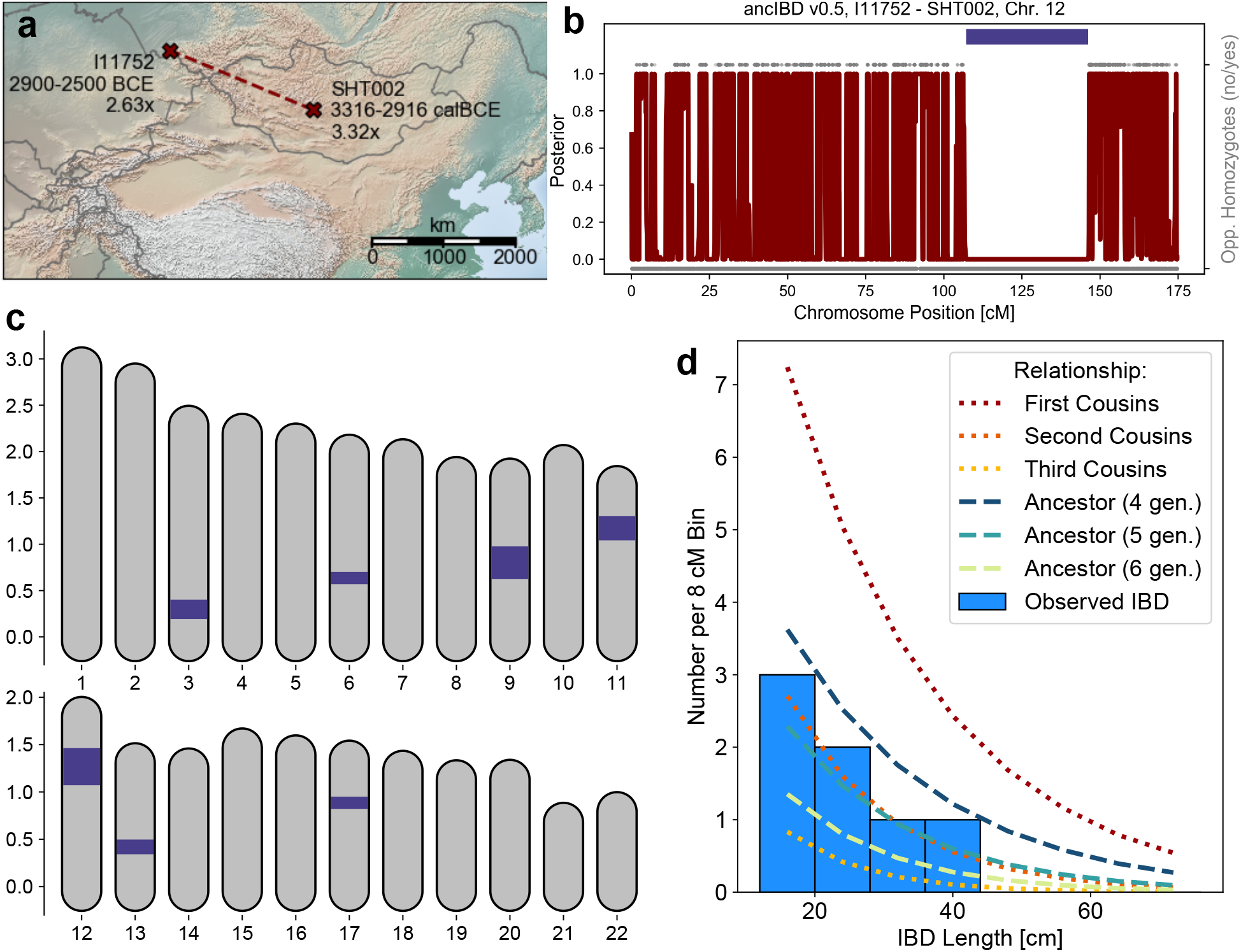
A geographically distant pair of ancient biological relatives detected with ancIBD. When screening ancient Eurasian individuals for IBD segments (see Fig. 3), we detected a pair of biological relatives whose remains were buried 1410 km apart, one in central Mongolia and one in Southern Russia. (Panel a). The two individuals were previously published in two different publications [Narasimhan et al., 2019, Jeong et al., 2020]. Both individuals are archaeologically associated with the Afanasievo culture and genetically cluster with other Afanasievo individuals [Narasimhan et al., 2019, Jeong et al., 2020]. **Panel b:** Posterior of non-IBD state on Chromosome 12, which has the longest inferred IBD segment (39.1 cM long, indicated as a dark blue bar). We also plot opposing homozygotes (gray dots up), whose absence is a necessary signal of IBD. Only SNPs where both markers have an imputed genotype probability greater than 0.99 are plotted. **Panel c:** Plot of all inferred IBD segments longer than 12 cM. **Panel d:** Histogram of inferred IBD segment lengths, as well as theoretical expectations for various types of relatives (calculated using formulas described in [Ringbauer et al., 2021]. Panels b-d were all created using default plotting functions bundled into the ancIBD software package.

Moreover, there are several intriguing observations regarding individuals associated with the Corded Ware culture, who are the earliest Central and Northern Europeans to carry high amounts of Steppe-like ancestry. Previous aDNA research has shown them to be a genetic mix of previous Final Neolithic farmer cultures as well as Steppe-like ancestry [Haak et al., 2015, Allentoft et al., 2015, Papac et al., 2021]. Using IBD, we find that individuals from diverse Corded Ware cultural groups, including from Sweden (associated with the Battle Axe culture), Russia (Fatyanovo), and East/Central Europe share high amounts of long IBD with each other, and also have IBD sharing up to 20 cM with various Yamnaya groups (Fig. 5, Fig. E2). Importantly, we find a distinctive IBD signal with the so-called Globular Amphora culture, in particular from Poland and Ukraine, who were Copper Age farmers not yet carrying Steppe-like ancestry [Mathieson et al., 2018, Schroeder et al., 2019]. This IBD link to Globular Amphora appears for all Corded Ware groups in our analysis, including from as far away as Scandinavia and Russia (Fig. 5), which indicates that individuals related to Globular Amphora contexts from Eastern Europe must have had a major demographic impact early on in the genetic admixtures giving rise to various Corded Ware groups.

**Figure 5:**
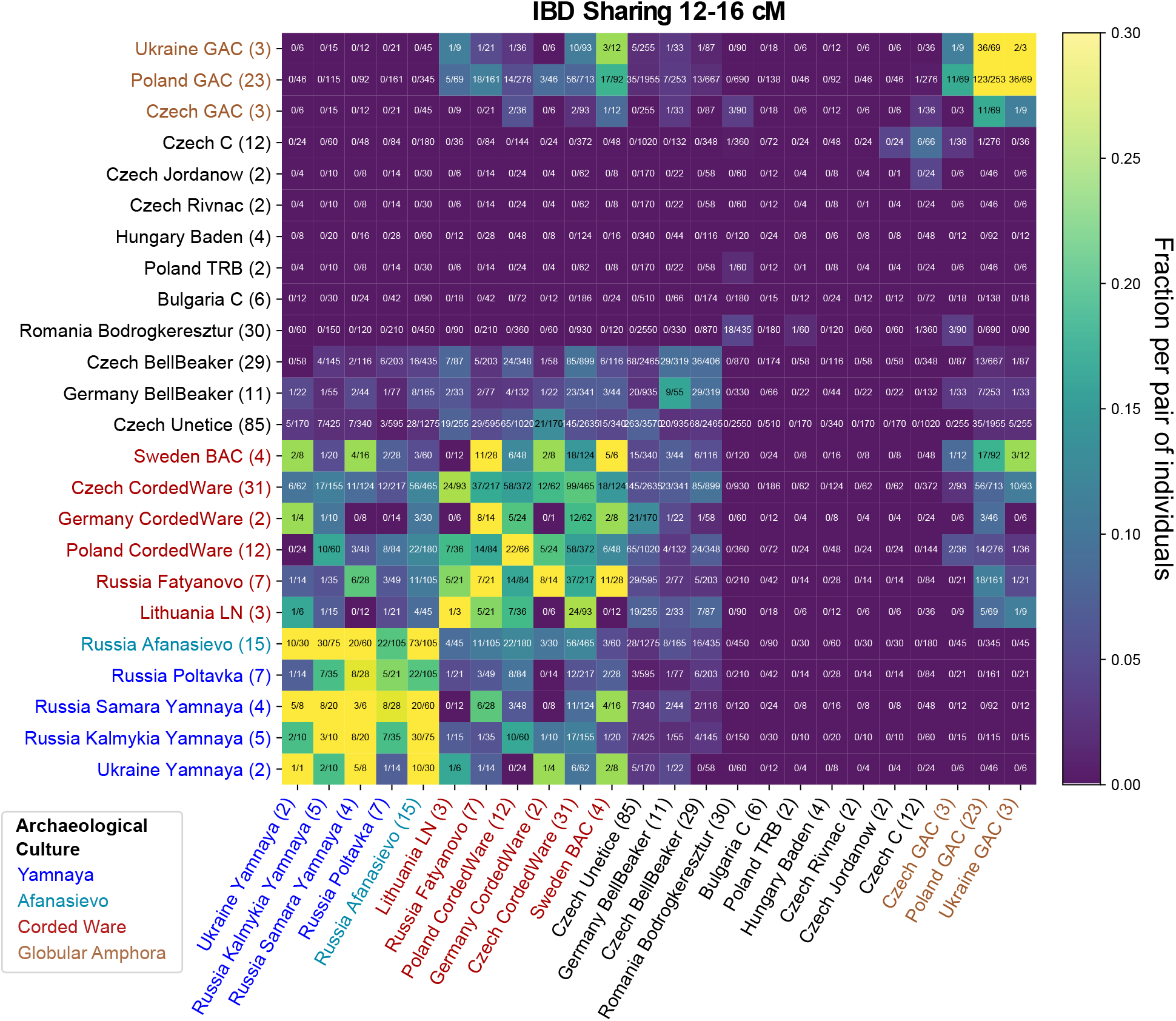
Inferred IBD segments between various Copper & Bronze Age West Eurasian Groups. We visualize IBD segments 12-16 centimorgan long (for IBD sharing in other length classes see Fig. E2). We applied ancIBD to identify IBD segments between all pairs of 304 West Eurasian ancient individuals (all previously published data, see Supp. Tab. 1c) organized into 24 Archaeological groups. For each pair of groups, we plot the fraction of all possible pairs of individuals that share at least one IBD 12-16 centimorgan long, which we obtained by dividing the total number of pairs that share such IBD segments by the total number of all possible pairs: Between two different groups of *n*_1_ and *n*_2_ individuals, one has *n*_1_ · *n*_2_ pairs, while within a group (on the diagonal in the figure) of size *n* one has *n*(*n* – 1)/2 pairs.

## Discussion

We have introduced ancIBD, a new method to detect IBD segments in aDNA data, and demonstrated its power to reveal close and distant biological relatives in systematic evaluations as well as empirical applications. Our algorithm follows a long line of work utilizing probabilistic HMMs to screen for IBD segments [e.g. Bercovici et al., 2010, Browning and Browning, 2013b, Vieira et al., 2016, Nait Saada et al., 2020, Severson et al., 2022b]. Our benchmarks have demonstrated that ancIBD robustly detects IBD longer than 8 centimorgan, for WGS data down to 0.25x and 1240k data down to 1x, and with WGS data performing generally better than 1240k data at the same average coverage depth. When compared to other methods to detect IBD, a particular advantage of ancIBD is its balanced performance on both precision and recall in the low coverage regime.

There are several extensions that could potentially improve the performance of ancIBD. We have not investigated how performance varies along the genome, but both SNP density in the 1240k and 1000G SNP set varies substantially along the genome [see e.g. Ringbauer et al., 2021], and thus likely also power and error rates when calling IBD. We have introduced a simple heuristic to filter out a small number of genomic regions with high false positive rates of long IBD. Focusing on regions of high SNP density could allow one to confidently call IBD with shorter lengths in specific parts of the genome. We also note that we have phased the data using a modern reference haplotype panel, which likely yields decreasing imputation and phasing performance the older the sample [Biddanda et al., 2022]. Future efforts to include ancient genomes into reference haplotype panels or to use modern reference panels substantially larger than 1000 Genomes will likely improve the quality of imputed ancient genomes, and thus also increase the performance of ancIBD.

Successful identification of long IBD extends the detection of biological relatives to up to the sixth-degree [Caballero et al., 2019], while allele-sharing-based methods are generally limited to detect relatives only up to the third-degree relatives [Lipatov et al., 2015, Monroy Kuhn et al., 2018]. A recent method KIN [Popli et al., 2023] fits transitions between IBD states to identify close relatives that share large fractions of their genome in IBD. While applicable to samples with lower WGS coverage (~ 0.05x) than ancIBD, this algorithm is limited by design to detecting biological relatives up to the third degree but is not tuned toward calling precise locations of sporadic IBD segments typical of more distant relatives.

A salient future application of detecting IBD in aDNA is improving the dating of ancient samples. Clearly indicating close relatedness, the sharing of long IBD between two individuals can be used to tether samples together in time and thus refine radiocarbon (C14) dates. In particular, such work can build upon existing Bayesian methods to refine C14 dates based on external information [Buck et al., 1991, Sedig et al., 2021, Massy et al., 2022].

Being able to identify IBD segments from intermediate coverage ancient DNA data unlocks a powerful new way to investigate fine-scale genealogical connections of past human populations. To showcase this potential, we have used ancIBD to generate new evidence regarding the origins of the people culturally associated with the Corded Ware Complex, who became the primary vector for the 3rd millennium BCE large-scale spread of steppe ancestry to Central and Western Europe [Allentoft et al., 2015, Haak et al., 2015, Olalde et al., 2018]. We show that the quarter of Corded Ware Complex ancestry which is specifically associated with earlier European farmers can be pinpointed distinctively via sharing of long IBD to people associated with the Globular Amphora culture of Eastern Europe, while the remaining three quarters must share recent co-ancestry with Yamnaya steppe pastoralists in the late third millennium BCE. This direct evidence that most Corded Ware ancestry must have genealogical links to people associated with Yamnaya culture spanning on the order of at most a few hundred years is clearly inconsistent with the hypothesis that the steppe ancestry in the Corded Ware primarily reflects an origin in as-of-now unsampled cultures genetically similar to the Yamnaya but in fact, related to them only a millennium earlier. The new information added by long haplotype sharing relative to average single-locus correlation statistics that have been the workhorse of aDNA studies to date lies in its ability to provide bounds on the number of generations separating pairs of individuals. A similar signal has recently been utilized by runs of homozygosity in aDNA [e.g. Racimo et al., 2020, Ringbauer et al., 2021], but now we are no longer restricted to haplotype sharing within one individual, but between all pairs of individuals.

Detecting IBD segments in modern DNA has yielded unprecedented insight into the recent demographic structure of present-day populations, allowing researchers to infer population size dynamics [Browning and Browning, 2015, Al-Asadi et al., 2019], genealogical connections between various groups of people [Ralph and Coop, 2013, Han et al., 2017, Nait Saada et al., 2020], and the geographic scale of individual mobility [Ringbauer et al., 2017, Al-Asadi et al., 2019]. In principle, such analysis can also be applied to ancient DNA. It is particularly promising that the number of sample pairs that can be screened for IBD grows quadratically with sample size. The rapidly growing aDNA record together with this even quicker growth of pairs that can be screened for IBD will provide aDNA researchers with a powerful new way to address demographic questions about the human past. We believe that the method to detect IBD in aDNA presented here is only a first step towards creating a new generation of demographic inference tools, giving insights into the human past at an unprecedented fine scale.

## Code Availability

A Python package implementing the method is available on the Python Package Index (https://pypi.org/project/ancIBD/) and can be installed via *pip*. Online documentation is available at https://ancibd.readthedocs.io/en/latest/index.html. Code developed for simulating data, analysis, and data visualization presented here is available at the GitHub repository https://github.com/hringbauer/ancIBD.

## Data Availability

No new DNA data were generated for this study. The reference panel data that we used for imputation (phased haplotypes from the 1000 Genomes dataset) are available at http://ftp.1000genomes.ebi.ac.uk/vol1/ftp/release/20130502/. The four high-coverage genomes used in empirical downsampling experiments were previously published [Wohns et al., 2022] and are available at https://reich.hms.harvard.edu/ancient-genome-diversit The AADR resource is publicly available at https://reich.hms.harvard.edu/allen-ancient-dna-resource-aadr-downloadable-genotypes-present-day-and-ancient-dna-data. The Hazleton samples can be downloaded through the European Nucleotide Archive under accession PRJEB46958. Raw sequencing data of the published Westeurasian ancient individuals are publicly available as documented in the original publications (see Supp. Tab. 1A).

## Author Contributions

We annotate author contributions using the CRediT Taxonomy labels (https://casrai.org/credit/). Where both authors serve in the same role, the degree of contribution is specified as ‘lead’, ‘equal’, or ‘support’.

- Conceptualization (Design of study) – lead: HR; support: DR, NP
- Software – HR; support: YH
- Formal Analysis – HR, YH
- Data Curation – DR, HR, AA
- Visualization – HR, YH
- Writing (original draft preparation) – HR, YH
- Writing (review and editing) – All authors
- Supervision – DR
- Funding Acquisition – DR

## Acknowledgement

We thank Shai Carmi (Hebrew University of Jerusalem) for his insightful comments on this manuscript. We gratefully acknowledge useful discussions with members of the Reich lab (Harvard University) and with the population genetics meeting group at the MPI-EVA Leipzig.

## Extended Figures

**Figure E1:**
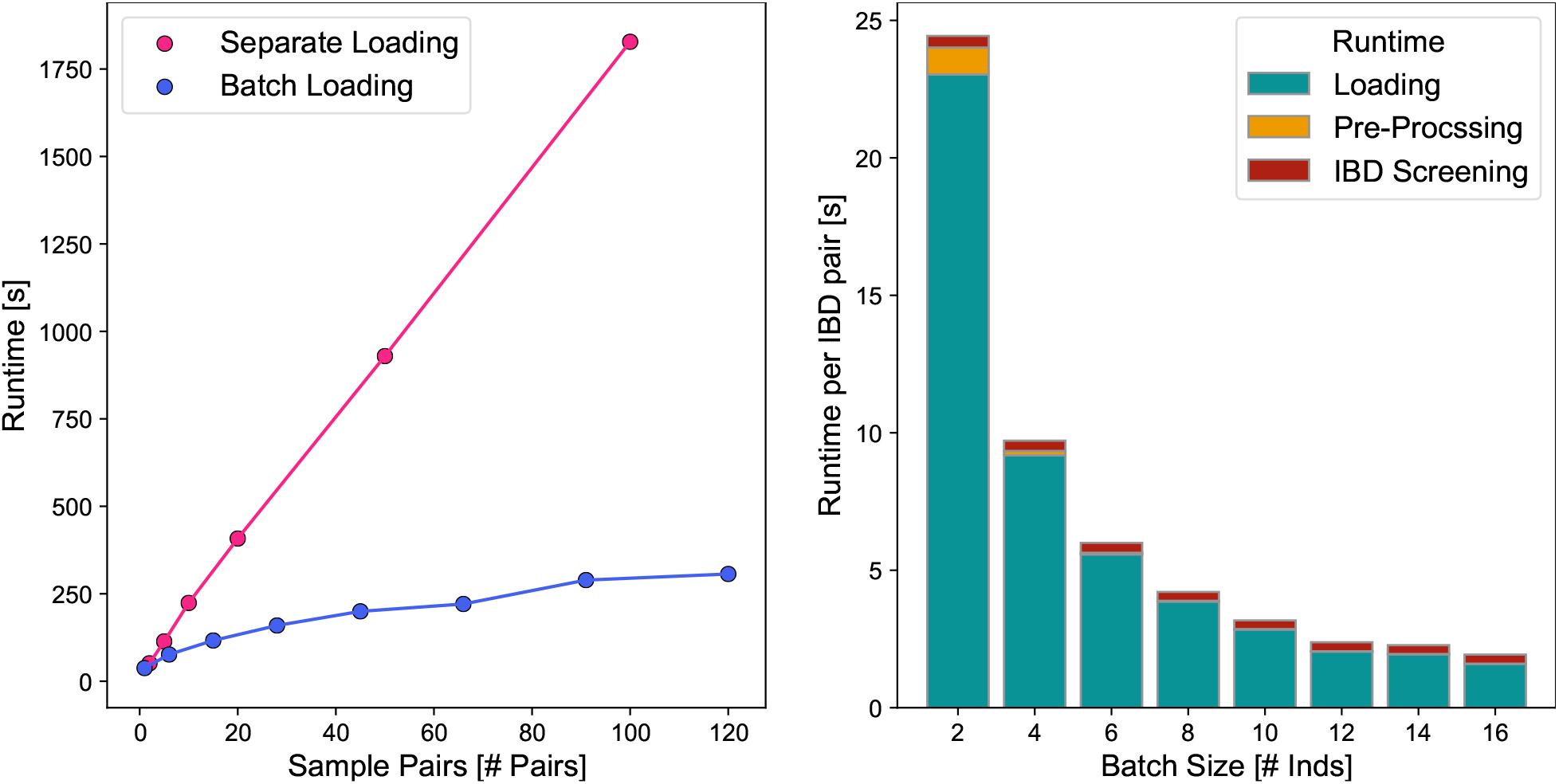
Run-time Benchmarks of ancIBD. To benchmark run-times, we applied ancIBD on empirical ancient DNA data in .hdf5 format imputed at 1240k sites. We used the imputed hdf5 file from the Eurasian application (Fig. 3), choosing samples and pairs at random. **Left:** For each sample pair, all autosomes are screened for IBD. In one experiment all pairs of samples were run independently, leading to a linear dependency on pair number, as expected. In a second experiment, all samples were loaded into memory, and then each sample pair was screened for IBD. The apparent sub-linear behavior is due to the fact that loading *n* samples scales slower than the actual runtime of *n*·(*n*–1)/2 sample pairs. **Right:** We depict the runtimes normalized per sample pair when screening all pairs of sample batches of various sizes for IBD. We visualize the loading time (the time it takes to load the hdf5 genotype data into memory), the pre-processing time (including preparing the transition and emission matrix), as well as the runtime of screening for IBD that includes the forward-backward algorithm as well as post-processing. Due to the decrease in the impact of the time to load the data, which scales linearly with batch size while the number of sample pair scales quadratically, we observe substantially increased runtimes per pair.

**Figure E2:**
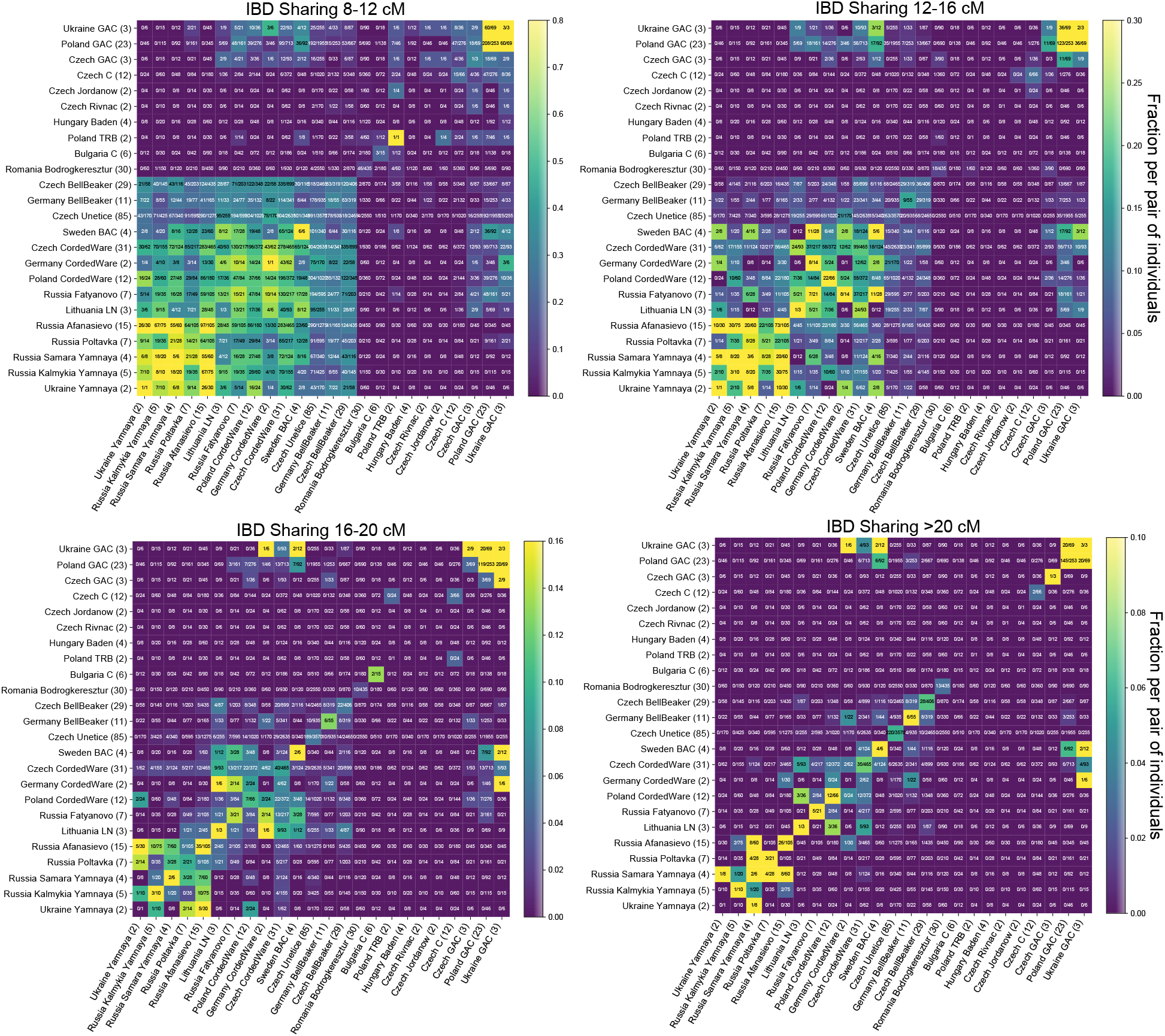
IBD sharing matrix of various Copper & Bronze Age West Eurasian Groups for four IBD length scales. As in Fig. 5, but for shared IBD [8 – 12 cM], [12 – 16 cM], [16 – 20 cM], > 20 cM long. We used ancIBD to infer IBD segments between all pairs of groups and visualize the fraction of pairs that share at least one IBD for each pair of populations and for the four different IBD length bins.

## Extended Tables

**Table E2:** Mean Inferred Segment Length for WGS-like Data

|  | 4cM | 8cM | 12cM | 16cM | 20cM |
| --- | --- | --- | --- | --- | --- |
| **2x** | 4.270 | 8.286 | 12.297 | 16.279 | 20.297 |
| **1x** | 4.341 | 8.351 | 12.334 | 16.311 | 20.349 |
| **0.75x** | 4.382 | 8.388 | 12.391 | 16.350 | 20.370 |
| **0.5x** | 4.448 | 8.469 | 12.454 | 16.422 | 20.431 |
| **0.25x** | 4.686 | 8.664 | 12.682 | 16.663 | 20.652 |
| **0.1x** | 5.182 | 9.657 | 13.755 | 17.957 | 21.606 |

**Table E3:** Mean Inferred Segment Length for 1240k-like Data

|  | 4cM | 8cM | 12cM | 16cM | 20cM |
| --- | --- | --- | --- | --- | --- |
| **2x** | 4.559 | 8.592 | 12.572 | 16.551 | 20.528 |
| **1x** | 4.825 | 8.961 | 12.956 | 17.000 | 20.860 |
| **0.75x** | 5.005 | 9.270 | 13.281 | 17.442 | 21.242 |
| **0.5x** | 5.196 | 10.003 | 14.287 | 18.391 | 22.094 |
| **0.25x** | 5.699 | 11.526 | 17.112 | 21.209 | 26.062 |
| **0.1x** | 6.017 | 12.002 | 19.161 | 25.150 | 30.370 |

