## Supplementary Notes for "ancIBD - Screening for identity by descent segments in human ancient DNA"

February 2023

| $\mathbf{g}$ | $P(\mathbf{g})$ | $P(\mathbf{g}\|D)$ | $P(\mathbf{g}\|s{=}1)$ |
|---|---|---|---|
| $(\mathbf{0},0,\mathbf{0},0)$ | $(1-p)^4$ | $(1-x_{1A})(1-x_{1B})(1-x_{2A})(1-x_{2B})$ | $(1-p)^3$ |
| $(\mathbf{0},1,\mathbf{0},0)$ | $p(1-p)^3$ | $(1-x_{1A})x_{1B}(1-x_{2A})(1-x_{2B}))$ | $p(1-p)^2$ |
| $(\mathbf{1},0,\mathbf{1},0)$ | $p^2(1-p)^2$ | $x_{1A}(1-x_{1B})x_{2A}(1-x_{2B})$ | $p(1-p)^2$ |
| $(\mathbf{1},1,\mathbf{1},0)$ | $p^3(1-p)$ | $x_{1A}x_{1B}x_{2A}(1-x_{2B})$ | $p^2(1-p)$ |
| $(\mathbf{0},0,\mathbf{0},1)$ | $p(1-p)^3$ | $(1-x_{1A})(1-x_{1B})(1-x_{2A})x_{2B}$ | $p(1-p)^2$ |
| $(\mathbf{0},1,\mathbf{0},1)$ | $p^2(1-p)^2$ | $(1-x_{1A})x_{1B}(1-x_{2A})x_{2B}$ | $p^2(1-p)$ |
| $(\mathbf{1},0,\mathbf{1},1)$ | $p^3(1-p)$ | $x_{1A}(1-x_{1B})x_{2A}x_{2B}$ | $p^2(1-p)$ |
| $(\mathbf{1},1,\mathbf{1},1)$ | $p^4$ | $x_{1A}x_{1B}x_{2A}x_{2B}$ | $p^3$ |

**Table S1:** List of ingredients for calculating emission probability for $s{=}1$

### Supp. Note S1 Emission Probabilities for IBD states

Here we compute the HMM emission probability $P(D|s)$ for the first IBD state ($s{=}1$). Due to symmetry, the emission probabilities for all other IBD states ($s{=}2,3,4$) can be calculated analogously by simple rearrangement.

First, we consider all possible combinations of phased genotypes that are compatible with $s{=}1$. The IBD state $s{=}1$ encodes haplotypes $1A,1B$ as being IBD, thus the alleles on haplotype $1A,1B$ have to match and be both reference or alternative. A total of $2\times2\times2{=}8$ genotype configurations are compatible with $s{=}1$. These eight configurations with their corresponding $P(\mathbf{g}),P(\mathbf{g}|D),P(\mathbf{g}|s{=}1)$ are listed in Tab. S1. The other eight configurations are not possible, and we have $P(\mathbf{g}|s{=}1){=}0$ for those.

Therefore, summing over all possible genotype combinations $\mathbf{g}\in\{0,1\}\times\{0,1\}\times\{0,1\}\times\{0,1\}$, we obtain now:

$$
\begin{aligned}
P(D|s{=}1) \sim \sum_{\mathbf{g}\in\mathcal{G}} \frac{P(\mathbf{g}|D)}{P(\mathbf{g})} P(\mathbf{g}|s{=}1) = \\
&\frac{(1-x_{1A})(1-x_{1B})(1-x_{2A})(1-x_{2B})}{1-p} + \frac{(1-x_{1A})x_{1B}(1-x_{2A})(1-x_{2B})}{1-p} \\
&+ \frac{x_{1A}(1-x_{1B})x_{2A}(1-x_{2B})}{p} + \frac{x_{1A}x_{1B}x_{2A}(1-x_{2B})}{p} \\
&+ \frac{(1-x_{1A})(1-x_{1B})(1-x_{2A})x_{2B}}{1-p} + \frac{(1-x_{1A})x_{1B}(1-x_{2A})x_{2B}}{1-p} \\
&+ \frac{x_{1A}(1-x_{1B})x_{2A}x_{2B}}{p} + \frac{x_{1A}x_{1B}x_{2A}x_{2B}}{p} \\
&= \frac{(1-x_{1A})(1-x_{2A})(1-x_{2B})}{1-p} + \frac{x_{1A}x_{2A}(1-x_{2B})}{p} + \frac{(1-x_{1A})(1-x_{2A})x_{2B}}{1-p} + \frac{x_{1A}x_{2A}x_{2B}}{p} \\
&= \frac{x_{1A}x_{2A}}{p} + \frac{(1-x_{1A})(1-x_{2A})}{1-p}
\end{aligned}
$$

### Supp. Note S2 Inserting IBD into Mosaic Simulations

We aimed to address two key issues when simulating IBD segments in aDNA data. First, we want to mimic errors and uncertainties of typical imputed aDNA data, such as those caused by low coverage and post-mortem damage. Second, we want to have accurate ground-truth IBD segments with exactly defined boundaries to be able to accurately assess length biases. Towards these two goals, we started from the 1000 Genomes Phase 3 release [Consortium et al., 2015] to simulate genotype data on the 1240k target sites of chromosome 3 in a two-step procedure. The first step establishes the ground-truth IBD segments. The second step captures the uncertainties of the imputation procedure.

In the first step, we simulated ground-truth genotypes by copying from haplotypes with the TSI group label (Tuscany, Italy) in the 1000 Genomes data, as has been done previously in Browning and Browning [2011], Ralph and Coop [2013]. We copied TSI haplotypes in blocks of 0.25 cM length, where each block was chosen randomly from all TSI samples. Any background IBD longer than 0.25 cM existing in the 1000 Genomes TSI samples is most likely broken up in such mosaic haplotypes, while fine-scale background LD patterns are mostly maintained. We then grouped pairs of haplotypes into diploid genomes. To add ground-truth IBD blocks, we overwrote one of the haplotypes of a pair of diploid samples. The start and end point of the overwrite was chosen randomly along the simulated chromosome, with the length of the overwrite matching specified segment lengths. Overall, we simulated IBD segments 4,8,12,16,20 cM long, each with 500 replicates.

In the second step, we use downsampled empirical aDNA data to mimic errors introduced in the imputation process. Generally, imputation accuracy at SNPs depends on allele frequencies, and homozygotes are better imputed than heterozygotes [e.g. da Mota et al., 2022]. To simulate these dependencies, we downsampled 52 high-coverage ancient samples (50 of them >15x average coverage, and two >10x, all double-stranded library and half-UDG treated) from AGDP (see data availability) to various target coverages. We determined ground-truth genotypes by imputing the original high-coverage data after clipping 5 base pairs from both ends of aligned sequencing reads to reduce aDNA damage. For each of the three possible genotypes (0/0, 0/1, 1/1) at each site, we assembled, for each coverage, a list of imputed genotypes and their associated genotype probabilities from the downsampled data. We then simulated imputation error at low coverage by setting the genotype and genotype probability at each SNP to those of a sample chosen randomly from the aforementioned list that matches the true genotype. In case a genotype is not found in any of the 52 high-coverage samples, we kept the true simu-

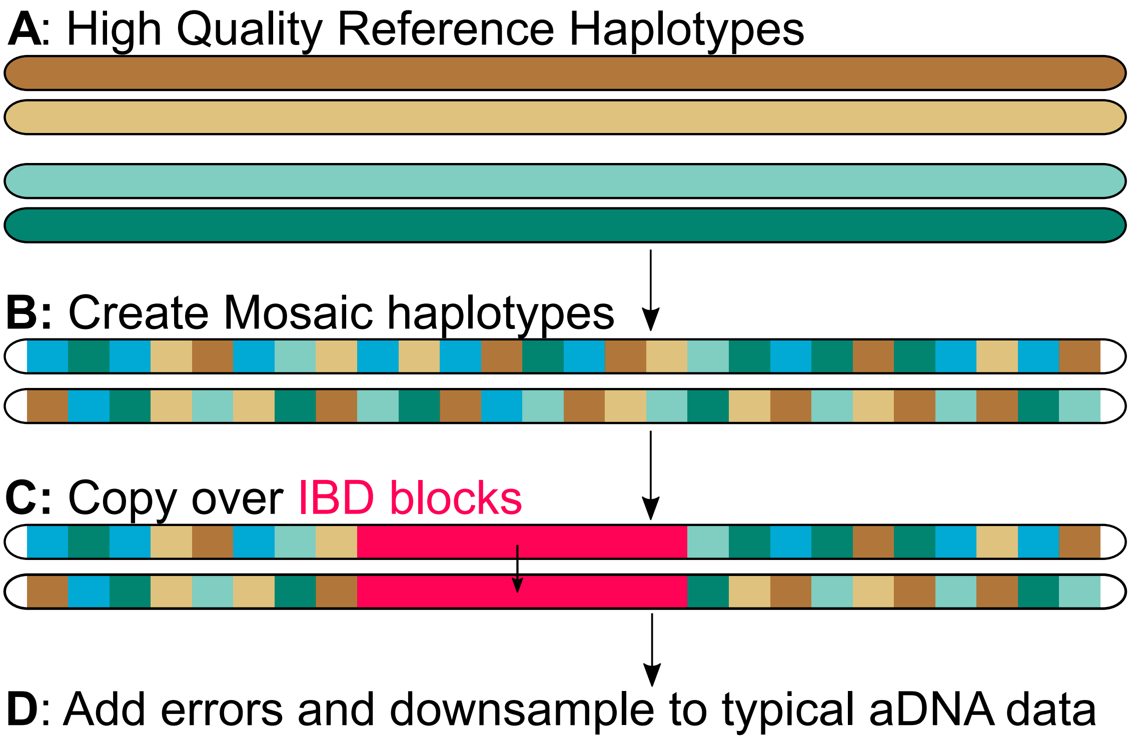


**Figure S1: Copying in IBD segments into haplotype mosaics.** We visualize our pipeline to simulate IBD segments starting from high-quality reference haplotypes as described in Supp. Note S2. We note that we grouped two mosaic haplotypes to obtain diploid individuals. To simplify visualization, we left out this second haplotype per individual from our sketch.

lated genotype and set its associated genotype probability to 99%. We note that of all 77,652 biallelic 1240k markers on chromosome 3, 52,629 have all three possible genotypes found in at least one of the 52 genomes, and 15,700 of them have at least two. While effectively sampling individual genotype probabilities from a mixture of downsampled samples, this procedure better mimics varying imputation accuracy at SNPs of various allele frequencies and genotype states [Hui et al., 2020].

Finally, we introduced phasing errors by flipping the phase at intervals drawn from an exponential distribution. To specify the mean of this distribution, we matched the average phased block length estimated from downsampling a high-quality trio (I3388, I3950, I3949, whose high-coverage WGS data were published in Wohns et al. [2022] and 1240k data in Narasimhan et al. [2019]). Both WGS and 1240k BAM files of this trio set were downsampled to 2x, 1x, 0.75x, 0.5x, 0.25x, 0.1x and then imputed and phased with GLIMPSE as described in Supp. Note S4. We identified phase switches between 1240k SNPs using VCFtools –diff-switch-error [Danecek et al., 2011]. The average phase block lengths for WGS and 1240k are summarized in Tab. S2.

| Coverage | WGS [cM] | 1240k [cM] |
|---|---|---|
| **2x** | 0.182 | 0.182 |
| **1x** | 0.257 | 0.125 |
| **0.75x** | 0.271 | 0.106 |
| **0.5x** | 0.271 | 0.0814 |
| **0.25x** | 0.220 | 0.0532 |
| **0.1x** | 0.127 | 0.0344 |

**Table S2: Mean Phased Block Length for WGS and 1240k Data at Various average Coverages.** All lengths are map lengths in centimorgan. Phase switch errors are inferred as described in Supp. Note S2 by downsampling a high-coverage ancient trio, using the high-coverage data as ground truth.

### Supp. Note S3 Imputation Pipeline

The imputation of aDNA is performed in the same way as in Waldman et al. [2022]. We first generated genotype probabilities using bcftools mpileup (v1.10.2) [Li, 2011] (with -q 30 -Q 30 filtering to use only high-quality aligned reads and bases). We imputed all autosomal bi-allelic SNPs 1000 Genomes Phase 3 release using GLIMPSE using its default parameters.

### Supp. Note S4 Downsampling Empirical aDNA Data

To assess the performance of ancIBD on realistic aDNA data, we downsampled high-coverage (∼20x) empirical human ancient DNA data. To obtain ground truth IBD segments to compare to, we used four WGS samples associated with the Afanasievo culture: I2105 (23.0X, 3300-2500 BCE, Ukraine, [Mathieson et al., 2018]); I3950 (25.8x, 2879-2632 calBCE, Russia, [Narasimhan et al., 2019]); I5273 (22.4x, 3011-2885 calBCE, [Narasimhan et al., 2019]) and I5279 (28.4x, 3011-2897 calBCE, Russia, [Narasimhan et al., 2019]).

To establish ground-truth diploid genotypes for those four samples, we computed genotype likelihoods from the high-coverage BAM files using bcftools and then applied GLIMPSE to impute diploid genotypes. We then filtered to transversion sites and called IBD segments with IBIS [Seidman et al., 2020]. This algorithm takes as input unphased diploid genotypes and utilizes the fact that in the absence of genotyping error, two samples cannot be homozygous for two different alleles ("opposing homozygotes") within an IBD region as the two samples have one of their haplotypes identical. This signal establishes a necessary condition for IBD, and the absence of opposing homozygotes over a long genomic region constitutes distinct evidence for IBD. The reason we chose IBIS to establish ground truth IBD segments is that very few high-coverage trio samples are available for aDNA and computational phasing with 1000 Genomes reference panel produces relatively high switch error rates (Tab. S2, [da Mota et al., 2022]). After restricting to transversion biallelic sites and applying a posterior $GP{>}99\%$ and $MAF{>}1\%$ (minor allele frequency) filters, we obtained 3,756,564 sites for IBD screening. IBIS identified a total of 157 IBD segments longer than 4 cM among the six pairs of the four samples. We visually inspected those detected IBD segments to confirm that they are depleted of opposing homozygotes and removed 23 of them that contained regions with very low SNP density, typically segments located over centromeres or on chromosome ends. We used the resulting 134 segments as ground-truth IBD blocks for benchmarks discussed in the following.

For the four Afanasievo samples, both WGS and 1240k capture data are available. We downsampled the WGS BAM files to 2x, 1x, 0.75x, 0.5x, 0.25x and 0.1x coverage and the 1240k BAM files to 2x, 1x, 0.75x and 0.5x, each with 50 replicates. We applied the same bcftools+GLIMPSE imputation pipeline as described in Supp. Note S3 and then ran ancIBD using its default parameters. We computed the precision and recall of ancIBD at various length bins and coverages when compared to the ground-truth IBD set described above. Similarly, we also screened the downsampled data with IBIS, using the same 1240k SNP set. Within a given map length bins of [5cM, 6cM), [6cM, 8cM), [8cM, 12cM), and >12cM, we calculated precision as the fraction of all inferred IBD segments that have at least 50% of their length covered by any true segment of any size and recall as the fraction of the total length of all true IBD segments that are at least 50% covered by inferred IBD segments of any size. Our results are summarized in Fig. S3.

Most notably, we found that for the same coverage, WGS data substantially outperforms 1240k data. Particularly, we found that 0.25x WGS data yields similar IBD calling accuracy as 1-2x 1240k data, both for ancIBD and IBIS.

For long IBD segments (>12cM) that are of particular interest for detecting relatives, ancIBD achieves both high precision and recall ($>90\%$) for all coverages tested here. Errors for segments in these length ranges remain negligible for most downstream analyses. We find that IBIS has substantially reduced power to identify IBD at lower coverages (<0.25x for WGS and <1x for 1240k), despite maintaining a consistently high precision over all coverages. For intermediate range segments (8-12cM), IBIS maintains relatively high precision (>90%) at all coverages tested while having reduced power at low coverages. ancIBD maintains high recall (~80%) at all coverages while having less than 80% precision at 0.5x for 1240k data. Overall, our results demonstrate that ancIBD yield accurate IBD calling (~90% or higher precision) at $>0.25$x WGS and >1x 1240k data.

Our results indicate that for studies using shorter IBD segments (6-8 cM), which are often a main signal for demographic inference, greater care should be taken as false positive rates and false negatives are not trivial anymore. The default SNP density filtered (as described in the method section of the main article) reduces the ancIBD's recall for these shorter segments (~65-70%). To improve the performance of ancIBD, we designed genomic masks that filter IBD in regions prone to false positive IBD segments due to low SNP density (Fig. S4). To identify regions with excessive IBD sharing, we computed the average IBD sharing rate (>6cM) among 10,156 Eurasian ancient individuals (same set as in Fig. 3) in genomic windows of size 0.5 cM. We then designated regions to be masked as those whose sharing rate exceeds three standard deviations from the genome-wide average IBD sharing. The start and end point of each masked region was determined by the first windows (on the left and right) whose sharing rate equals or falls below the genome-wide average. With the mask applied, the precision of ancIBD without SNP density filtering remains as high as the one without mask and with SNP density filtering; however, we observe a substantial boost in power within the unmasked region to greater than $90\%$ (Fig. S5).

We note that the precision of ancIBD reported in those downsampling experiments should be interpreted as being conservative because we likely underestimate precision in our downsampling experiments for the following two reasons. First, our benchmarks indicate that IBIS prioritizes precision over recall, especially for shorter segments, as reported previously (Seidman et al. [2020], Fig. 3 and Fig. S4). Thus, IBIS might miss some true IBD segments in the high-coverage data that are called by ancIBD in the downsampled data. Second, we visually screened all the detected IBD segments and as ground truth only retained those that are depleted of opposing homozygotes without major gaps, which might effectively remove some true IBD segments.

To assess whether some IBD inferred by ancIBD are missing in the ground truth data set, we computed the rate of ”opposing homozygotes” for each detected segment using the genotypes called from the high-coverage BAM files. We define the rate of opposing homozygotes as the percentage of sites where two samples carry homozygotes for different alleles out of all sites where both samples carry homozygotes. We included only transversion sites with minor allele frequency $>10\%$ in the 1000G reference panel in this calculation so that the probability of being homozygote for both reference and alternative alleles is non-negligible. We then plotted this rate of opposing homozygotes against a segment's Positive Predictive Value (PPV), defined as the fraction of a called segment covered by any segments in the ground-truth set. We found many segments with low PPV that have rates of opposing homozygotes similarly low as segments with very high PPV (Fig. S6, S7, S8, S9). This observation indicates that in the ground-truth set least parts of true IBD segments are missed, which would decrease the precision of ancIBD. That said, it is hard to determine whether these segments of exceptionally low opposing homozygote rates are fully true IBD segments. Thus we chose to be conservative in our tests.

#### Supp. Note S5 Estimating False Positive Rates with Downsampled Empirical Data

False positive IBD segments are particularly problematic for many downstream analyses such as demographic inference; therefore, it is important to establish for which coverage and IBD length cutoffs the false positive rate is tolerable for a particular application. To estimate false positive rates from empirical data, we selected 13 ancient individuals (I4893, I4596, I1583, I2978, I5838, I1507, I2861, I2520, I3758, I5077, I0708, I5233, I3123) from AGDP (see Data Availability and Supp. Tab. 1F) that have both high-coverage WGS and 1240k aDNA data available. All samples are chosen to be from Western Eurasia so that their imputation quality is expected to be relatively homogenous and the estimated false positive rates are not driven by a subset of them being poorly imputed. We determined the ground-truth diploid genotypes on chromosome 3 as described in Supp. Note S4 and then used IBIS to confirm that these samples share no IBD with each other. We further verified the absence of IBD sharing by plotting opposing homozygous along chromosome 3 for visual inspection. Therefore, all inferred IBD segments from the downsampled data are false positives.

For WGS data, we downsampled to 0.1x, 0.25x, 0.5x, and 0.75x. For 1240k data, we downsampled to 1x, 2x, and 3x (for 3x downsampling, we excluded samples I5233 and I5077 due to insufficient coverage of the original 1240k BAM file). For each target coverage, we created 50 independent replicates and the estimated average false positive rates are visualized in Fig. 2b.

As in Supp. Note S4, we found that WGS data outperforms 1240k aDNA data of the same coverage. We also note that, depending on different applications, the coverage cutoff for ancIBD is different. For example, for detecting biological relatives using IBD segments longer than 12cM, a more lenient coverage requirement can be employed (0.25x for WGS data and 1x for 1240k data). For demographic inference, one must take into account the signal-to-noise ratio; therefore, the IBD length and coverage cutoff are dependent on the effective population size of the study population and should be decided on a case-by-case basis. In Fig. 2b we plotted expected IBD sharing for constant populations with different effective population sizes to aid such comparison. To calculate this expected sharing, we use established formulas for expected IBD in panmictic populations of constant size (see e.g. [Ringbauer et al., 2021, Fernandes et al., 2021]).

#### Supp. Note S6 Simulating IBD-sharing of biological relatives with PEDSIM

To gain insight into the number and length distribution of IBD blocks given degree of parental relatedness, one can calculate expected numbers of blocks falling into certain length classes [Ringbauer et al., 2021, see e.g.]. However, these calculations do not address the natural biological variance of the IBD distribution and also rely on the assumption that recombination can be modeled as a Poisson process (i.e. measuring genomic distances in Morgan) and do not incorporate the biological process of recombination interference (i.e. recombination events are less clustered than expected) as well as sex-specific recombination maps. While for relatives beyond second degree, these model violations have only little impact, these processes can significantly influence IBD patterns when an individual's parents are close relatives [Caballero et al., 2019].

For these reasons, we utilized the recently developed method ped-sim (v1.0.6) to simulate shared IBD segments between close relatives [Caballero et al., 2019]. For each degree of a given relatedness up to the sixth degree, we simulated $100$ pairs of individuals each, using the sex-specific genomic map of [Bhérer et al., 2017], and simulating all autosomes with the recombination interference model [Campbell et al., 2015] incorporated into PEDSIM. We visualize the simulated IBD sharing in Fig. 3b. We note that we simulated both ancestral relationships (e.g. parents and grand-parents) and also relationships via full sibs (e.g. full sibs themselves or uncle /aunts). These two relationships types can have different distributions of IBD lengths for the same degree of relatedness because the number of meiosis in relationships via full sibs is elevated by one while the number of shared haplotype ancestors is four instead of two.

#### Supp. Note S7 Comparison with other methods

In this section, we compare ancIBD's performance with other IBD callers designed for modern DNA data. To our knowledge, no dedicated IBD caller has been developed for ancient DNA previously; however, the same fundamental principles of detecting IBD segments apply to ancient and modern DNA. Thus, methods designed for modern DNA data might extend to imputed low-coverage aDNA data.

#### Supp. Note S7.1: IBDseq

IBDseq [Browning and Browning, 2013] is designed for whole genome sequencing data. It computes the likelihood ratio of IBD and non-IBD for each biallelic marker and then sums them to find long stretches of IBD regions. For applying IBDseq to imputed data, we filtered imputed variants to MAF >1% and imputation INFO score $>0.8$ (the same filtering as performed in Allentoft et al. [2022]). We merged the four ancient samples with 503 diploid samples from the 1000 Genome Project labeled as belonging to the super population EUR because IBDseq relies on population allele frequencies estimated from input samples. We found that for long segments (>12cM) and high coverage, both ancIBD and IBDseq perform equally well. In addition, compared with ancIBD, IBDseq has higher power in detecting intermediate segments (8-12cM) at higher coverage. However, IBDseq's precision quickly drops below an acceptable level for low coverages that are typical for most aDNA data (Fig.S10), especially for 1240k data. Additionally, we also tried to further filter imputed variants to only keep transversion sites to mitigate the effect of aDNA damage, however, we found that this filtering has only negligible effects on IBDseq's precision and recall (Fig.S11).

#### Supp. Note S7.2: GERMLINE and GERMLINE 2

Both GERMLINE [Gusev et al., 2009] and GERMLINE 2 [Nait Saada et al., 2020] rely on accurate phasing as they take a seed-and-extend approach to search for identical haplotypes between two samples. For GERMLINE, we used the same SNP filtering as for IBDseq described above. We attempted to tune default parameters to accommodate the noisy nature of imputed aDNA data (e.g, turn on the '-g_extend' option recommended for noisy data, allowing up to 10 mismatch homozygous and heterozygous markers per slice); however, we could not identify any setting that enabled GERMLINE or GERMLINE 2 to detect any IBD segments among the test samples. Having effectively zero power is most likely due to the relatively high switch error rates in aDNA data imputed with modern reference panels (Tab. S2), which is an order of magnitude higher than what is attained for modern DNA phased with biobank scale reference panel [da Mota et al., 2022, Rubinacci et al., 2021].

##### Supp. Note S7.3: hapIBD

Similar to GERMLINE, hapIBD [Zhou et al., 2020] requires phased genotypes. We used the same SNP filtering as for IBDseq and adjusted hapIBD's default parameters to allow more mismatches (min-seed=0.1, min-extend=0.05, max-gap=500000, where the default values for the three parameters are 2.0, 1.0, 1000, respectively). Despite those attempts, hapIBD's power remains very low and the detected segments tend to be highly fragmented (Fig.S12), making it generally not applicable for imputed aDNA data.

#### Supp. Note S8 Other Supplementary Figures

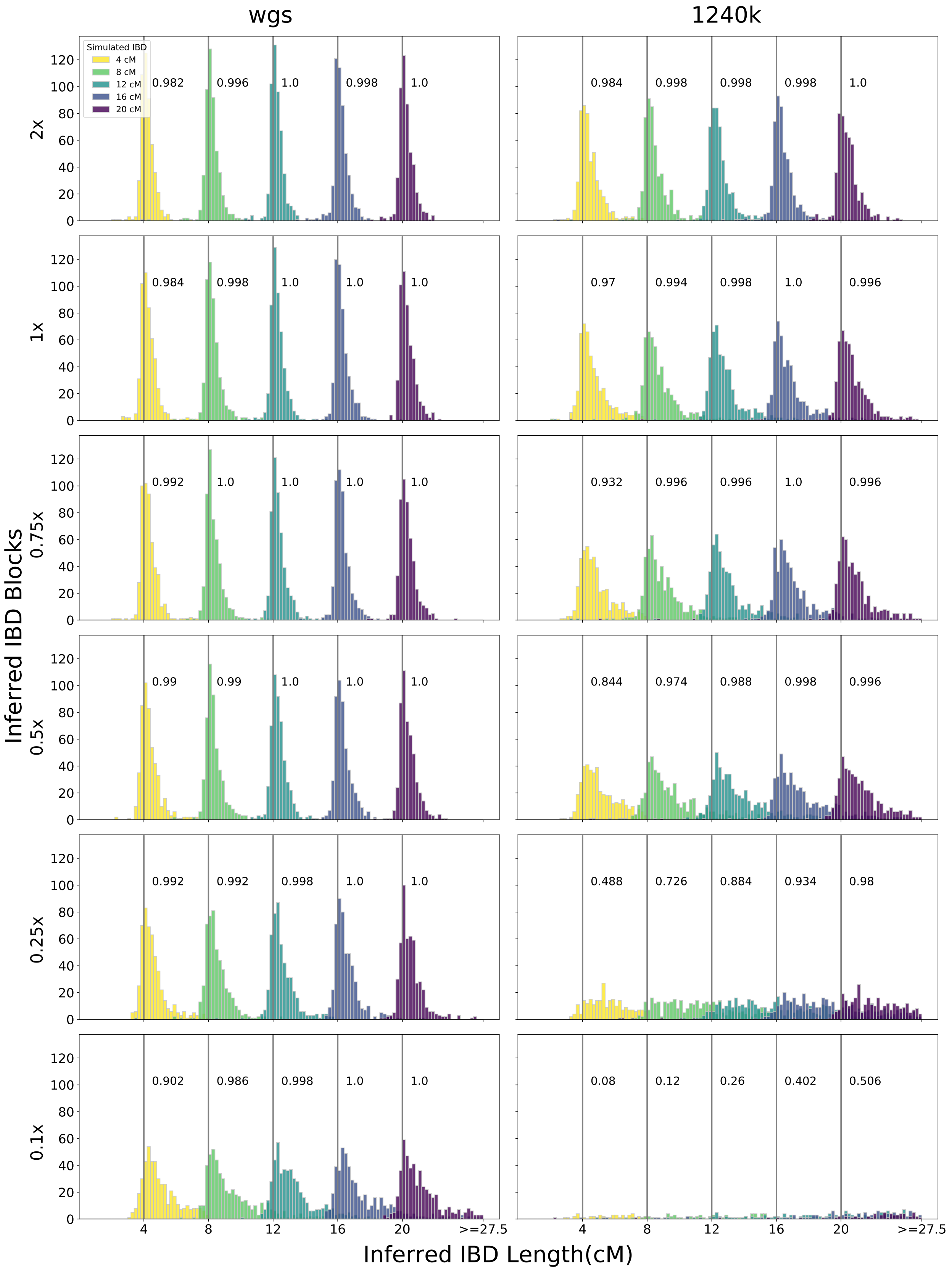


**Figure S2: IBD Calling Accuracy.** Accuracy of IBD calling in simulated synthetic diploid samples with IBD segments of length 4,8,12,16,20 cM. We simulated shotgun-like and 1240k-like data as described in Supp. Supp. Note S2). We visualize false positive, power, and general length bias for coverages from 2x down to 0.1x (rows). We also show false positive IBD segments longer than 4cM (red) and indicate power to call segments of each simulated length next to the respective gray vertical bars.

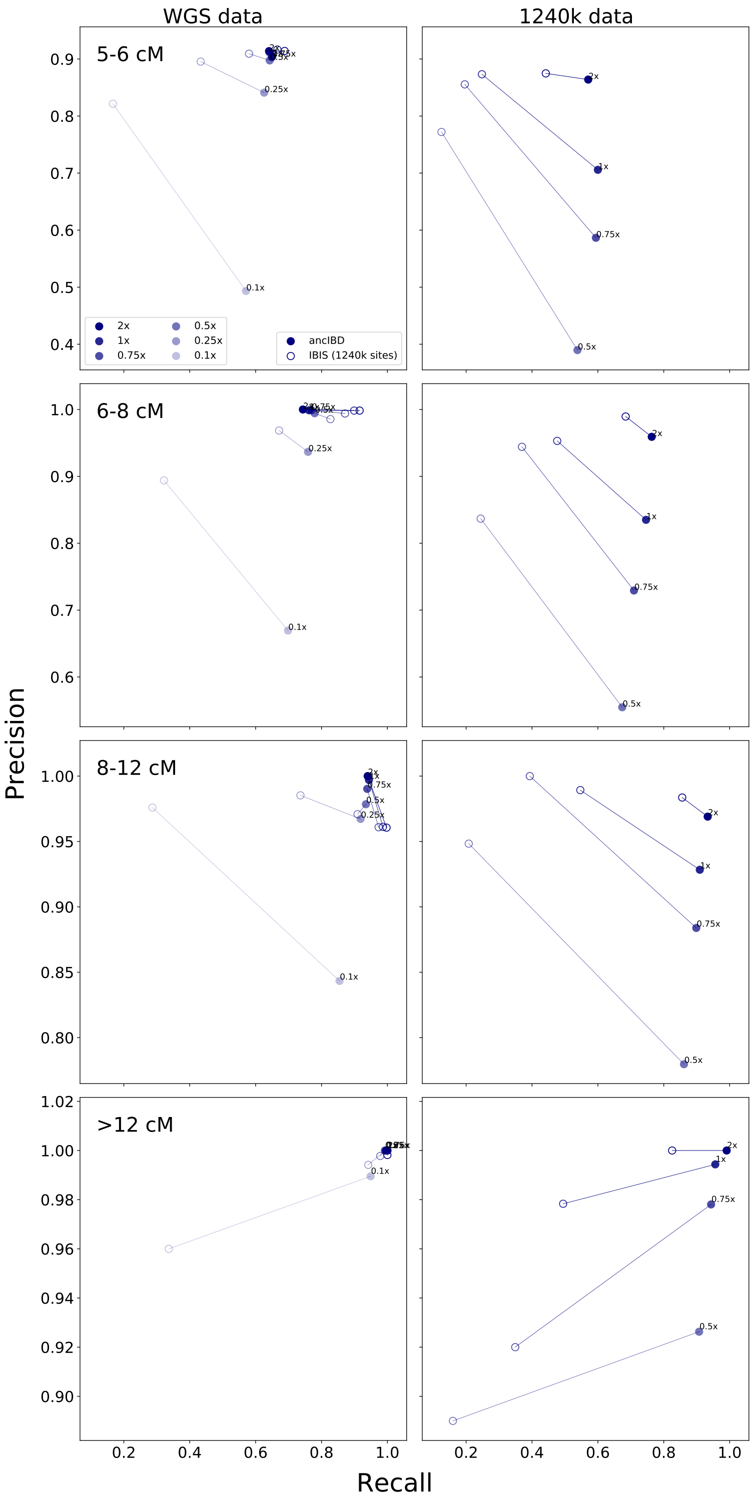


**Figure S3: Precision and recall of ancIBD and IBIS at various length bins and coverages.** We applied both methods with their default settings to genotype data imputed after downsampling to various coverages. For each coverage, we report the average precision and recall of each length bin across 50 independent replicates. Each row represents a length bin and each column represents one input data type (either WGS data or 1240k data). Note that the y-axis ranges are different for different rows.

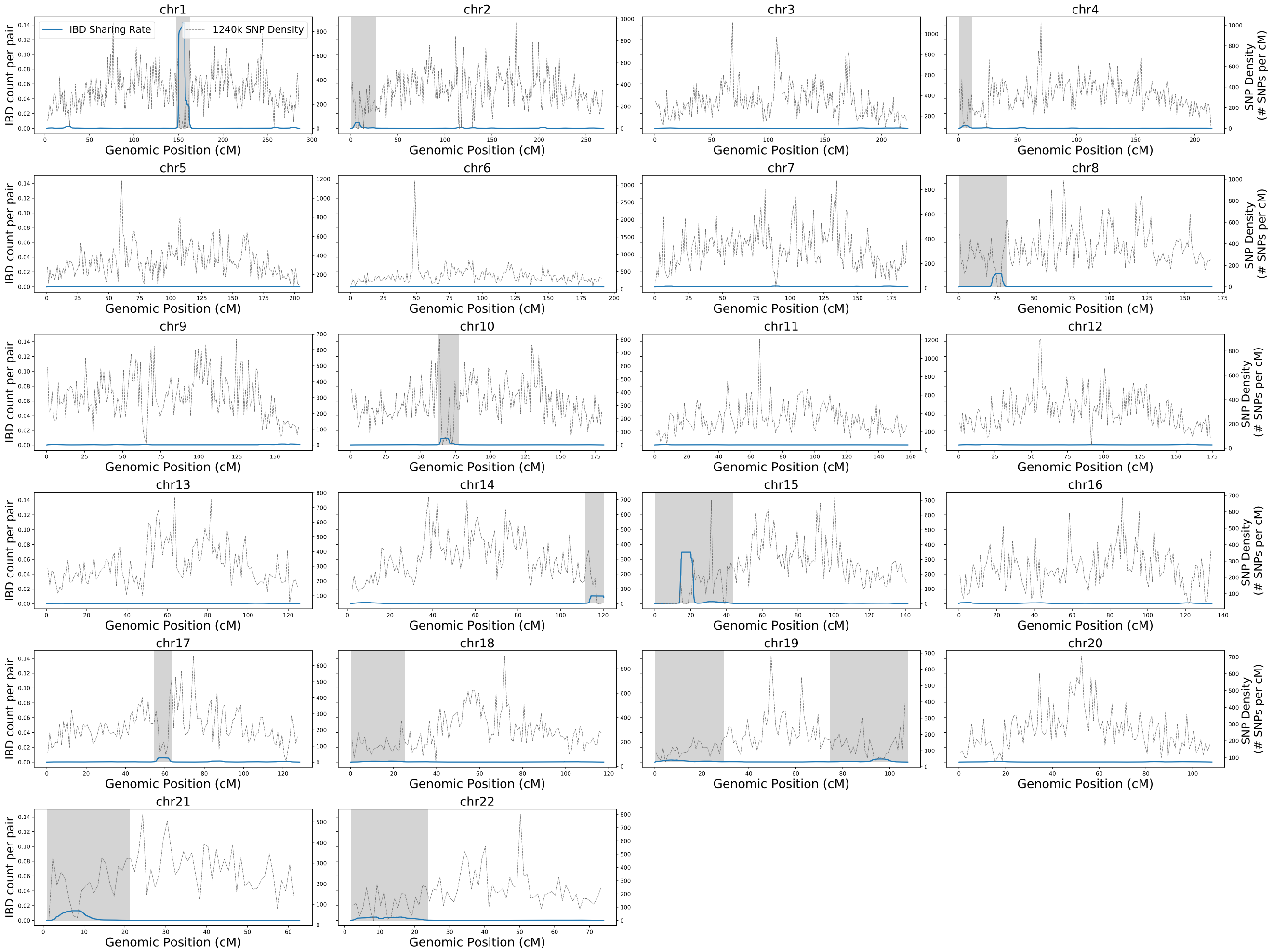


**Figure S4: IBD Sharing Rates along the genome.** Average genome-wide IBD sharing rate on the 22 autosomes plotted for all the 1240k target sites. We indicate regions with excessive sharing of IBD that are excluded when using our mask (gray areas). The average sharing rate was computed from the IBD inferred between 10,156 ancient individuals described in the main manuscript.

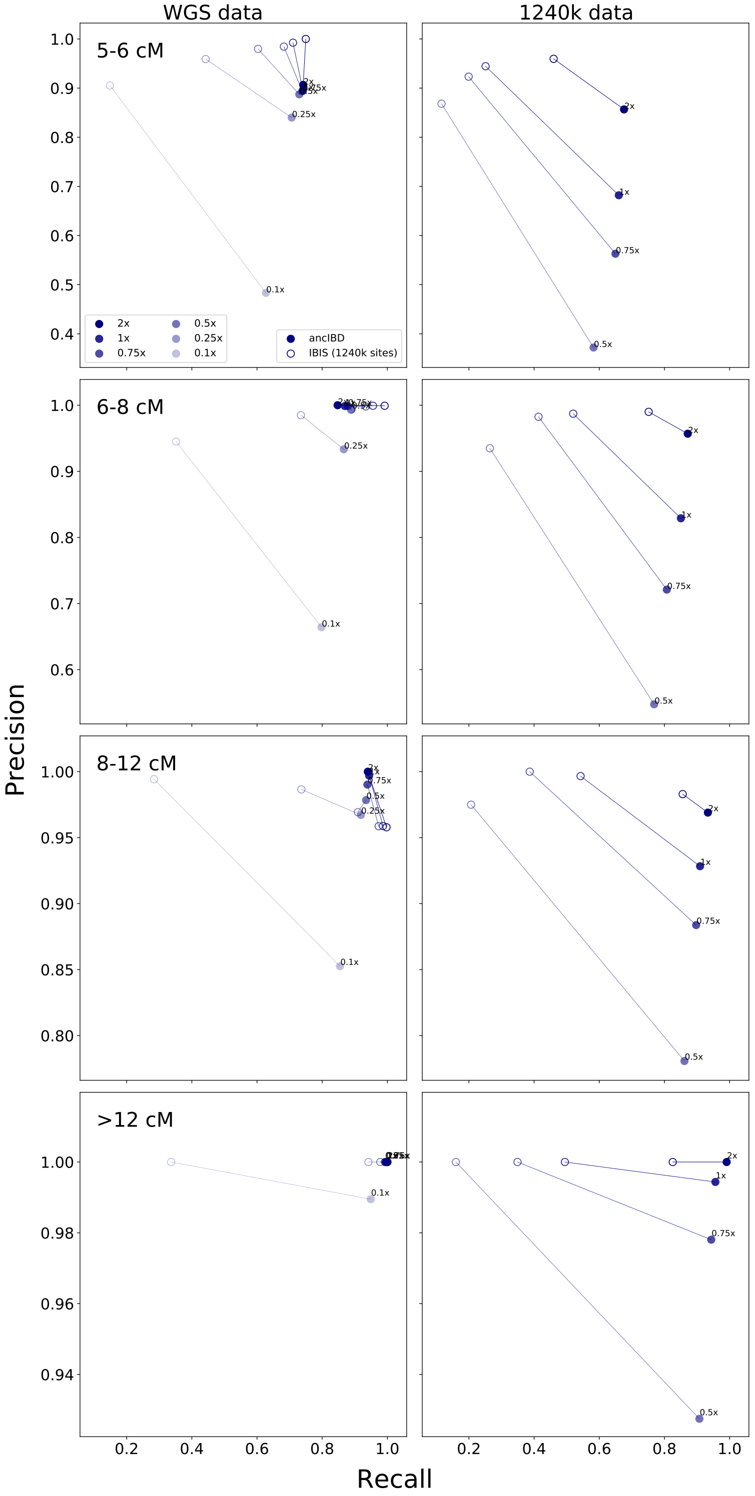


**Figure S5: Precision and recall of ancIBD and IBIS for various IBD length bins and coverages.** We applied ancIBD using the genomic masks as shown in Fig.S4 and without SNP density filtering. All other settings are the same as in Fig. S3.

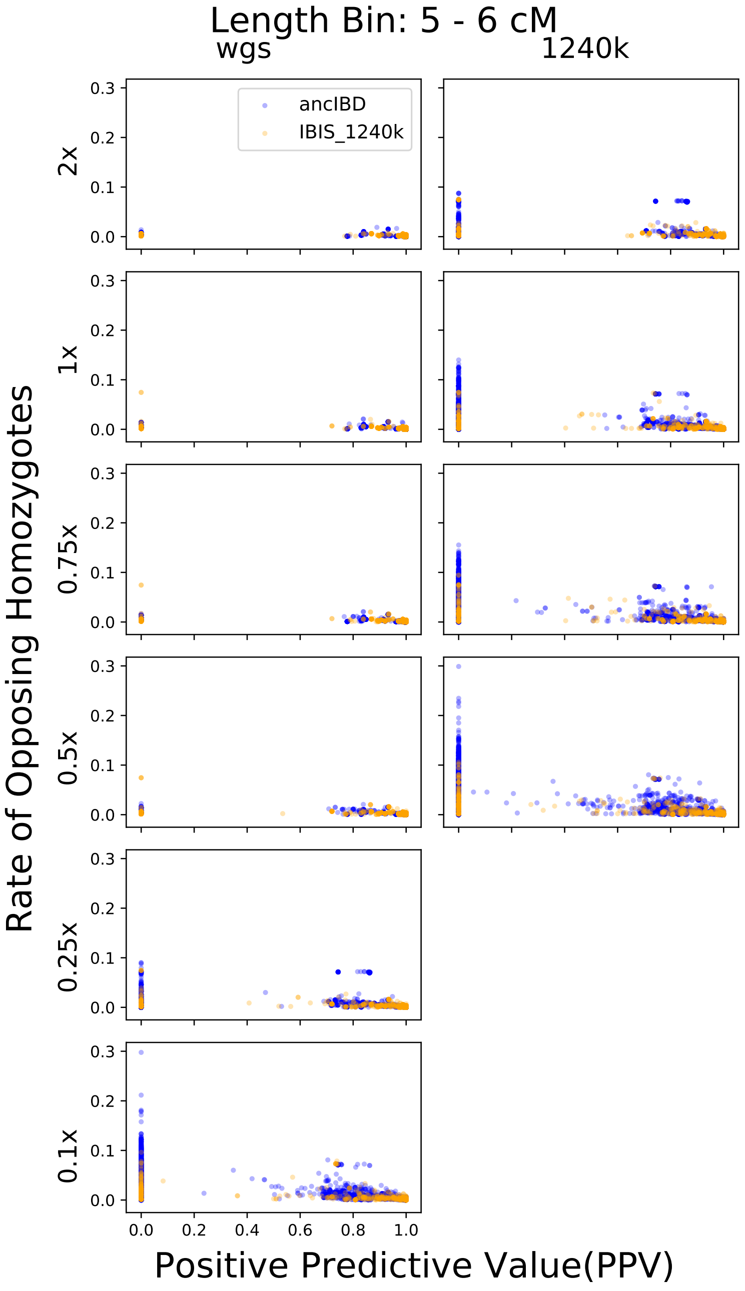


**Figure S6: Rate of Opposing Homozygotes for called segments in length bin 5-6cM.**

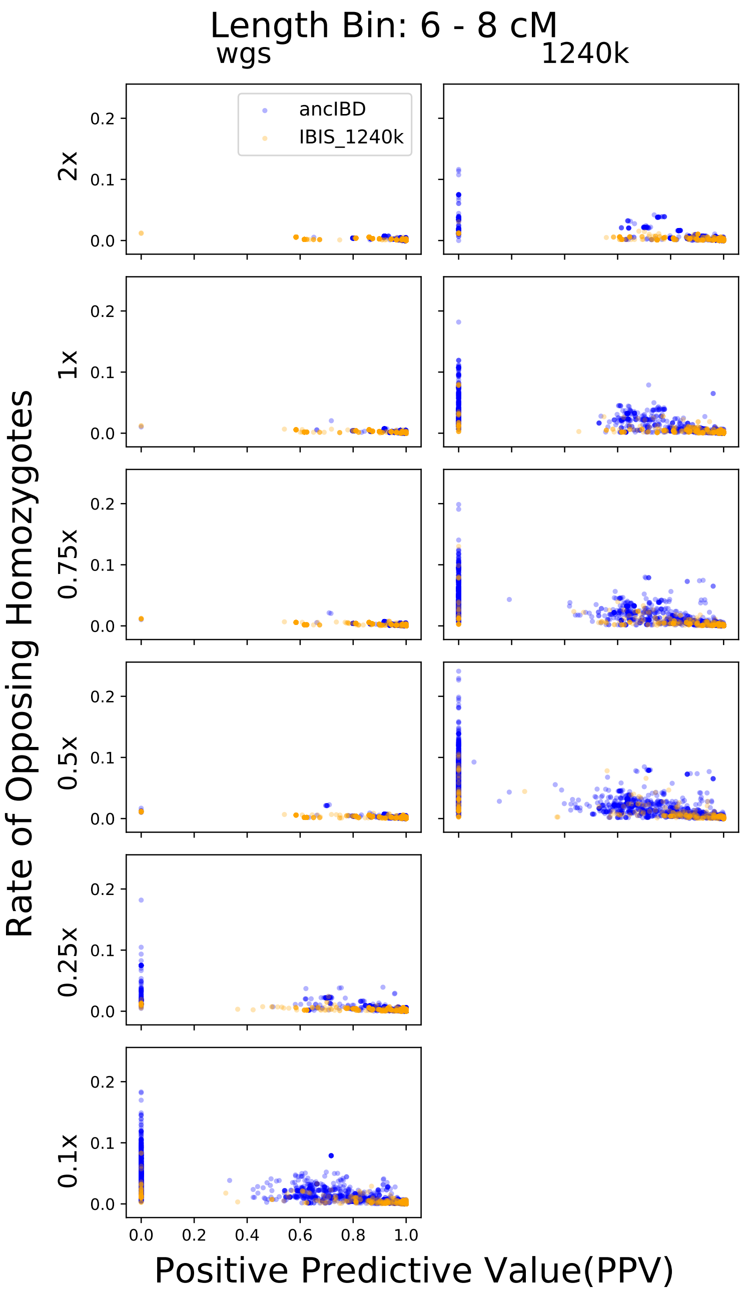


**Figure S7: Rate of Opposing Homozygotes for called segments in length bin 6-8cM.**

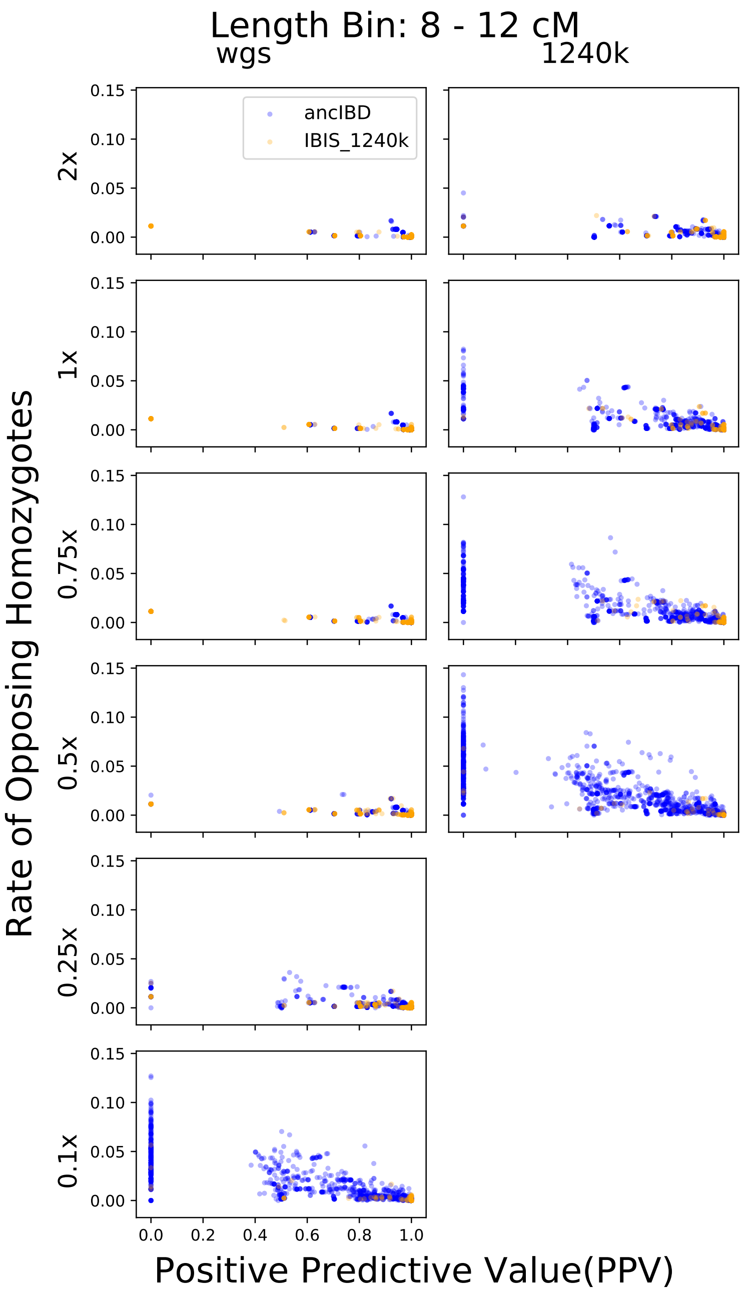


**Figure S8: Rate of Opposing Homozygotes for called segments in length bin 8-12cM.**

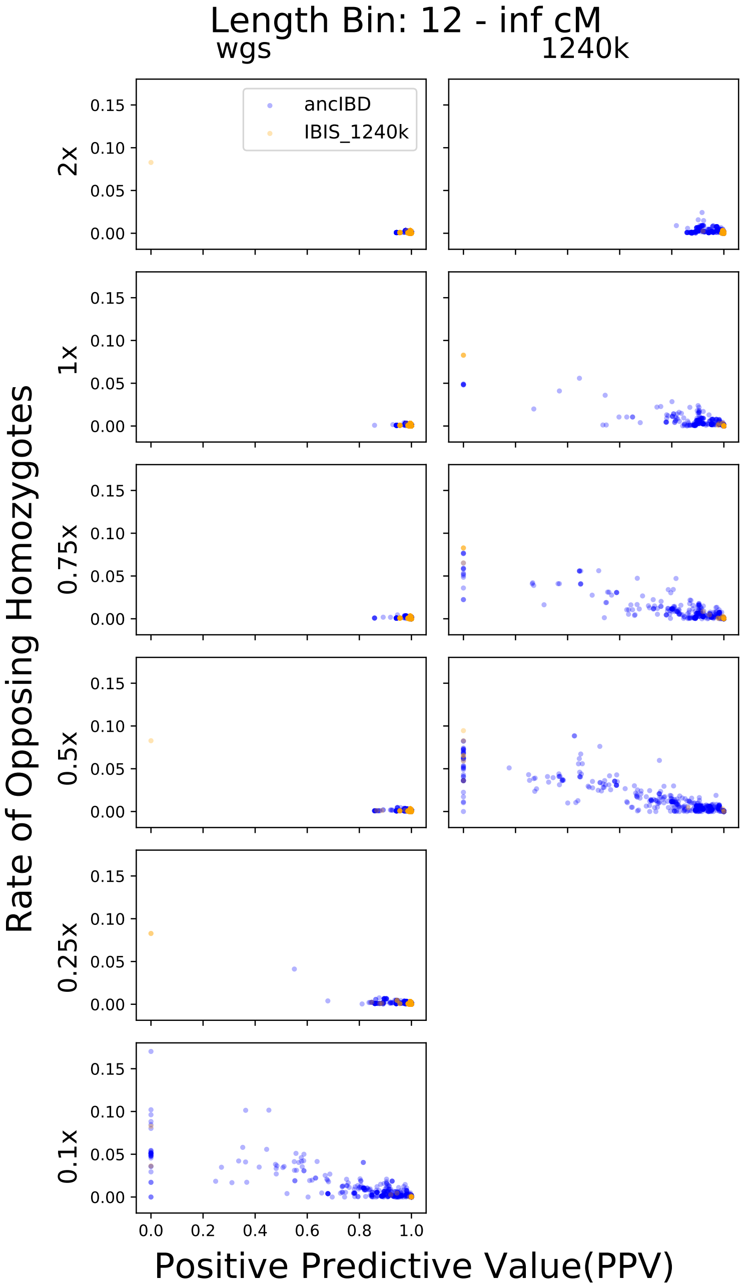


**Figure S9: Rate of Opposing Homozygotes for called segments in length bin ⩾12cM.**

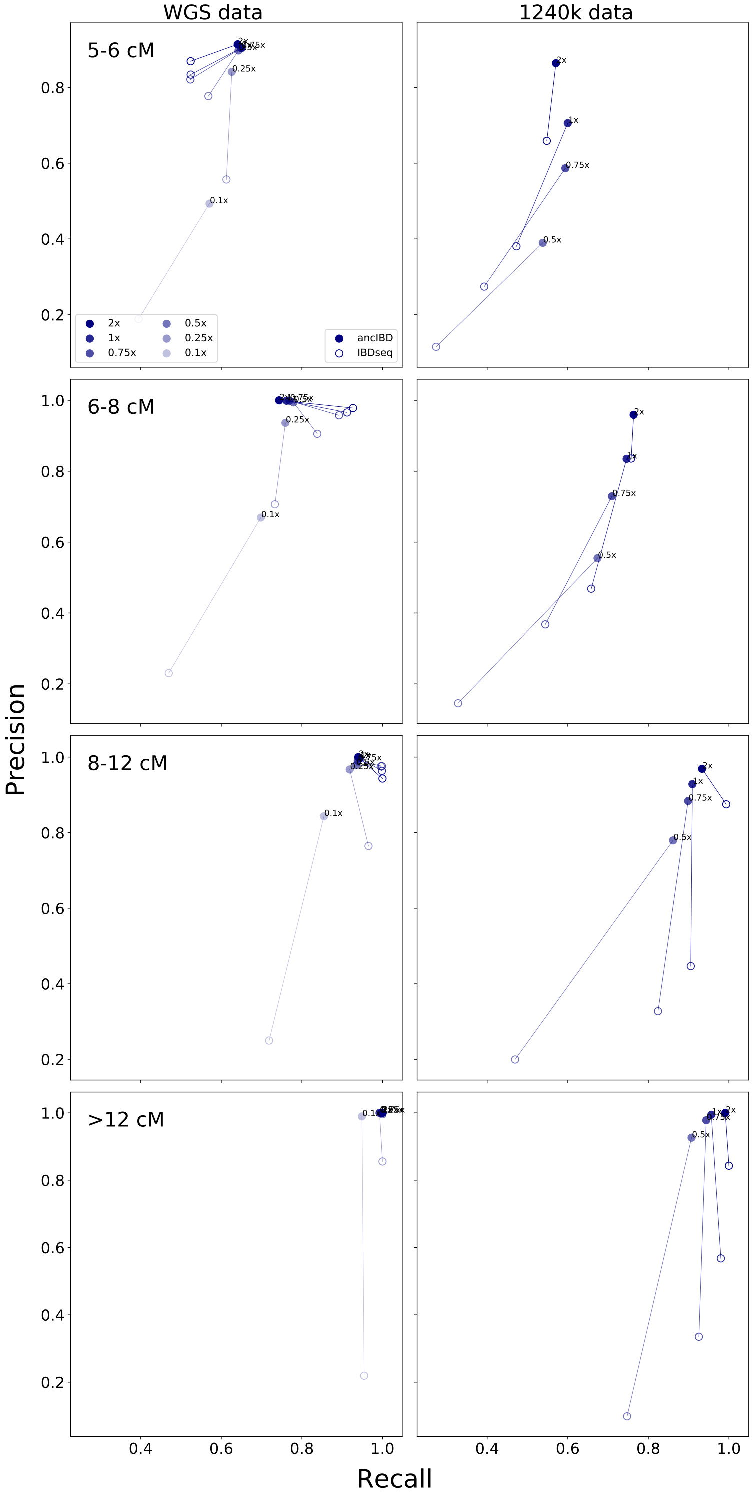


**Figure S10: Precision and recall of ancIBD and IBDseq at various length bins and coverages.** We applied IBDseq as described above and compared its precision and recall with ancIBD at various coverages and IBD length bins.

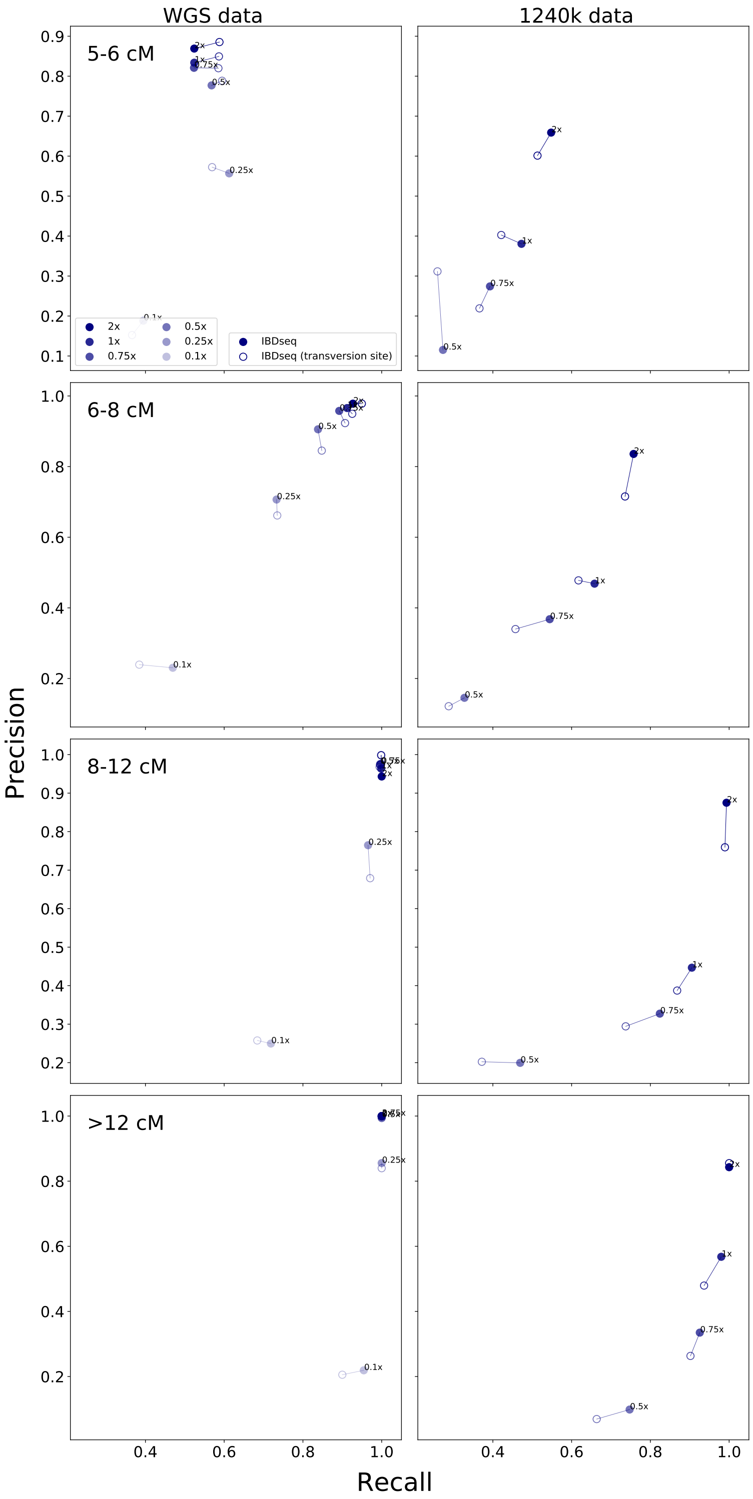


**Figure S11: Precision and recall of IBDseq with and without filtering transition SNPs.**

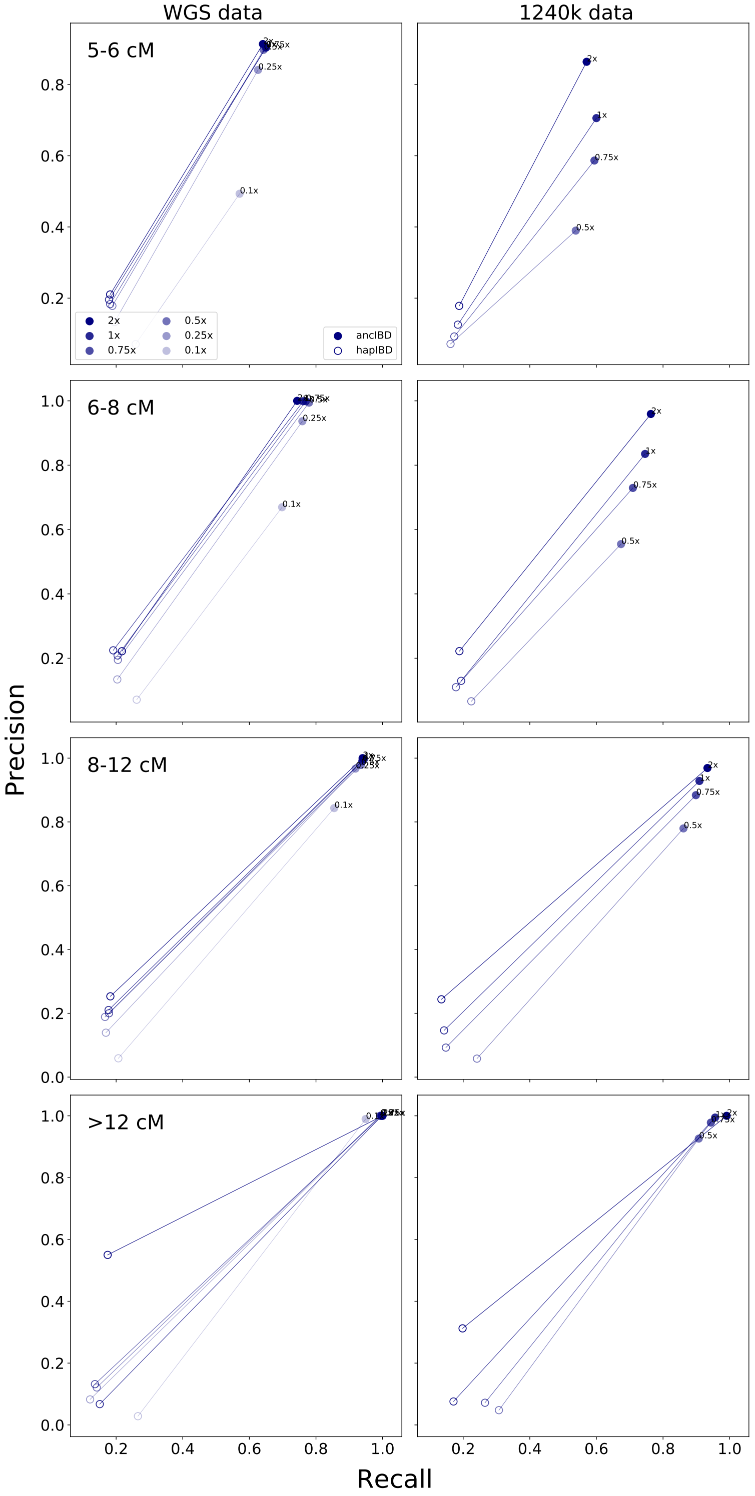


**Figure S12: Precision and recall of hapIBD at various length bins and coverages.** We applied hapIBD as described above at various coverages and computed its precision and recall for various IBD length bins.

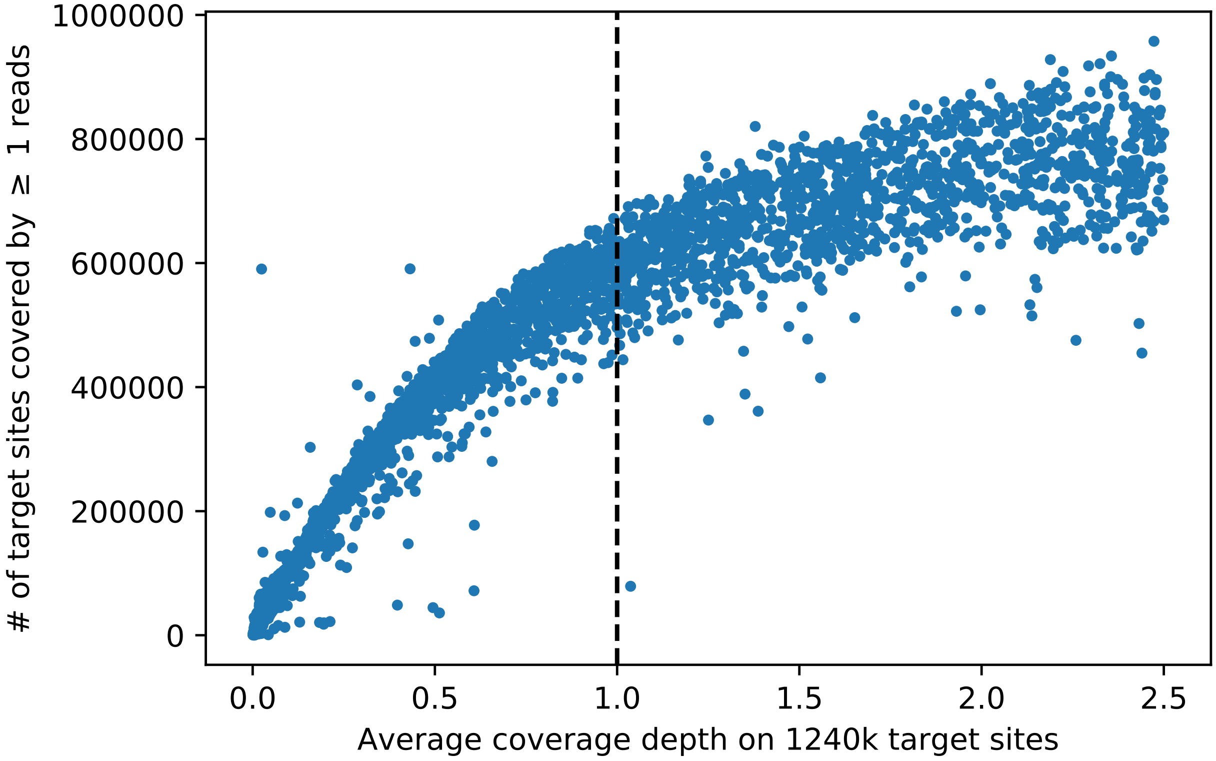


**Figure S13: Relationship between number of sites covered by ⩾ 1 reads and average coverage depth on 1240k SNP sites.** The plot shows the average coverage depth and the number of sites covered for 1240k samples from AADR (release v54.1). The recommended coverage cutoff (1x) is indicated by a black vertical dashed line. Only samples with less than 2.5x coverage are depicted.

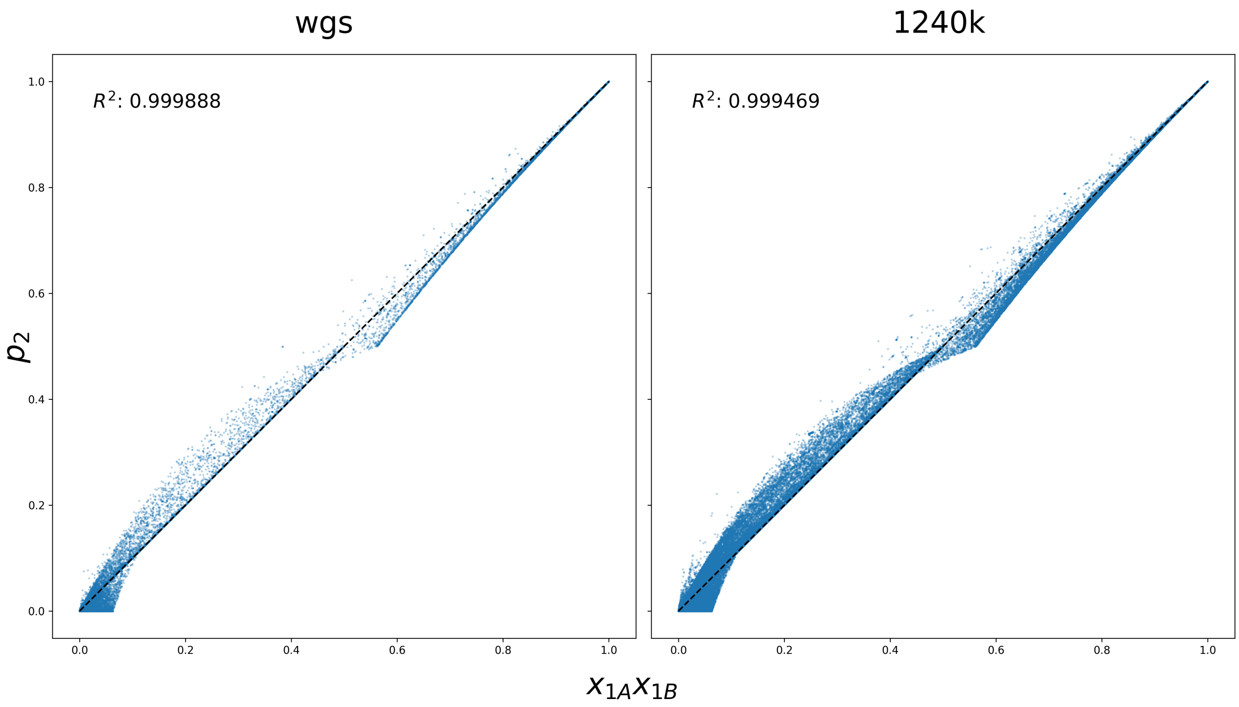


**Figure S14: Validity of approximating diploid genotype probabilities as the product of haplotype probabilities.** As described in the main text, we approximate $P(\mathbf{g}|D)$ as the product of the four probabilities of each of the haplotypes (1A, 1B) and (2A, 2B) being reference or alternative. Here we check the validity of this approximation by plotting $p_2$ against $x_{1A}x_{1B}$, where $p_2$ is the GLIMPSE-estimated genotype probability of being homozygous alternative alleles. The data points come from all the variants on chr1 in the 1000 Genome reference panel. The figure shows the result of I2105 downsampled to 1x. The coefficient of determination (calculated from sklearn.metrics.r2_score) is indicated in the upper left corner.
